# Restriction of *Wolbachia* bacteria in early embryogenesis of neotropical *Drosophila* species via ER-mediated autophagy

**DOI:** 10.1101/2021.04.23.441134

**Authors:** Anton Strunov, Katy Schmidt, Martin Kapun, Wolfgang J. Miller

**Affiliations:** Center for Anatomy and Cell Biology, Department of Cell and Developmental Biology, Medical University of Vienna, Austria; Department of Evolutionary Biology and Environmental Studies, University of Zurich, Switzerland

**Keywords:** Symbiosis, *Drosophila*, *Wolbachia*, tropism, autophagy, development, germline

## Abstract

*Wolbachia* bacteria are maternally transmitted intracellular microbes that are not only restricted to the reproductive organs but also found in various somatic tissues of their native hosts. The abundance of the endosymbiont in somatic tissues, usually a dead end for vertically transmitted bacteria, causes a multitude of effects on life history traits of their hosts, which are still not well understood. Thus, deciphering the host-symbiont interactions on a cellular level throughout a host’s lifecycle is of great importance to understand their homeostatic nature, persistence and spreading success. Using fluorescent and transmission electron microscopy, we conducted a comprehensive analysis of *Wolbachia* tropism in somatic and reproductive tissues of six *Drosophila* species at the intracellular level during host development. Our data uncovered diagnostic patterns of infections to embryonic primordial germ cells and to particular cells of somatic tissues in three different neotropical *Drosophila* species of the willistoni and saltans groups that have apparently evolved in both independently. We further found that restricted patterns of *Wolbachia* tropism are already determined in early fly embryogenesis. This is achieved via selective autophagy, and the restriction of infection is preserved through larval hatching and metamorphosis. We further uncovered tight interactions of *Wolbachia* with membranes of the endoplasmic reticulum, which might play a scaffolding role for autophagosome formation and subsequent elimination of the endosymbiont. Finally, by analyzing *D. simulans* lines transinfected with non-native *Wolbachia*, we uncovered that the host genetic background regulates tissue tropism of infection. Our data demonstrate a peculiar and novel mechanism to limit and spatially restrict bacterial infection in somatic tissues during a very early stage of host development.

## Introduction

*Wolbachia* are endosymbiotic bacteria residing within cells of many arthropod and some nematode species (reviewed in Kaur et al., 2021). Most of these host-microbe associations are considered facultative and even pathogenic (Min and Benzer, 1997), although cases of obligate mutualism also exist (Dedeine et al., 2003; Taylor et al., 2005; Hosokawa et al., 2010; Miller et al., 2010; Schneider et al., 2019). In insects, high trans-generational infectivity and maintenance of *Wolbachia* is ensured by its successful transovarial transmission (reviewed in Werren et al., 2008; Landmann, 2019), albeit cases of horizontal transmission also exist (reviewed in Pietri et al., 2016; Chrostek et al., 2017). Thus, the microbe mostly relies on colonization of the female germline to be stably transmitted to the next generation (Serbus et al., 2008; Kaur et al., 2021). However, the infection is not solely confined to reproductive organs and can be found in different somatic tissues like the central nervous system (CNS), retina, fat body, muscles, hemolymph and Malpighian tubules of a host (reviewed in Pietri et al., 2016). Such a variety of bacterial localization brings about a wide range of effects on host fitness and behavior (reviewed in Zug and Hammerstein, 2015). Moreover, regulation of *Wolbachia* density within somatic tissues is a key factor in host-symbiont association, strongly affecting both host survival and persistence of bacteria in a population (Min and Benzer 1997; Chrostek et al., 2013; Martinez et al., 2014; López-Madrigal and Duarte, 2019). The rich somatic life of the bacteria provides a scarcely studied repertoire of intimate cell-specific interactions balancing host-microbe association. Understanding its essence is of great importance for fundamental knowledge as well as for application in biological control of invertebrate pests and vectors of diseases (reviewed in Ross et al., 2019).

The neotropical *Drosophila* species *D. paulistorum*, *D. willistoni* and *D. tropicalis* (willistoni group) as well as *D. septentriosaltans* and *D. sturtevanti* (saltans group) represent unique models for studying host-microbe interactions due to their long-term history of co-evolution with *Wolbachia* endosymbionts (Miller and Riegler, 2006; Miller et al., 2010). Each of these neotropical *Drosophila* species carries a specific *Wolbachia* strain, which exhibits either obligate mutualistic (*D. paulistorum*) or facultative (all other four host species) relationships. Among these neotropical *Wolbachia* strains, *w*Pau, *w*Wil, *w*Tro and *w*Spt from *D. paulistorum*, *D. willistoni*, *D. tropicalis* and *D. septentriosaltans* are closely related to each other, whereas *w*Stv from *D. sturtevanti* is the most distantly related to the rest (Miller and Riegler, 2006; Martinez et al., 2014). In embryos of *D. willistoni* (Miller and Riegler, 2006) and *D. paulistorum* (Miller et al., 2010) native *Wolbachia* are mainly restricted to the primordial germ cells (PGCs), the future germline, whereas palearctic fly hosts like *D. melanogaster* and *D. simulans* embryos show systemic infections with no defined tropism (Miller and Riegler, 2006).

We have furthermore recently uncovered the spatial and asymmetric restriction of *Wolbachia* in *D. paulistorum* to defined larval and adult brain regions (Strunov et al., 2017), which might be linked to the symbiont-directed assortative mating behavior observed in this obligate host-microbe association (Miller et al., 2010; Schneider et al., 2019). However, it remains unclear if the PGC and neural restrictions are unique to *D. paulistorum* hosts, (*ii*) at which developmental stages the tropism is established and (*iii*) by which cellular mechanism(s) the germline and somatic *Wolbachia* restrictions are achieved. Such diverse types of host-microbe interactions provide an opportunity to decipher the mechanistic basis for their tropism to defined somatic and germline tissues as well as their density within a cell.

By using fluorescent *in situ* hybridization (FISH) with *Wolbachia*-specific probes throughout host development we uncovered spatial and temporal dynamics of both the “systemic” and “restricted” infection types in six native *Drosophila* hosts. With the help of sequential *Wolbachia*-FISH and immunofluorescence we showed that the distribution of infection is determined already during early embryogenesis with elimination of *Wolbachia* from most of the embryonic cells, but not PGCs, through autophagy. This leads to a restriction of infection to the future gonads and a few particular areas of somatic tissues in the adult. With the help of transmission electron microscopy, we mapped out the early stages of the bacterial elimination process and could demonstrate that the endoplasmic reticulum tightly encircling *Wolbachia* in early-cellularized blastodermal embryos might serve as a scaffold for assembly of the autophagy machinery. Finally, by transferring a natively restricted *Wolbachia* strain into a systemic background, we decipher that the host background plays a major role in regulating the infection tropism in tissues.

## Results

### *Wolbachia* infection is restricted to specific areas of somatic and reproductive tissues of some neotropical *Drosophila* species

In a recent publication, we have shown that, contrary to the systemic infections in *D. melanogaster* and *D. simulans* (Albertson et al., 2013), *Wolbachia* of neotropical *D. paulistorum* flies are tightly restricted to certain brain areas (Strunov et al., 2017). In the present study we investigated whether such an explicit isolation of infection in the nervous tissue is an exceptional case for *D. paulistorum* flies or similar examples of bacterial restriction could be found in other related species. We analyzed the distribution of native *Wolbachia* in both somatic and reproductive tissues of five neotropical *Drosophila* species (*D. paulistorum, D. willistoni*, *D. tropicalis*, *D. septentriosaltans*, *D. sturtevanti*) and *D. melanogaster* as a representative for the systemic infection (Strunov et al., 2017). Finally, we tested bacterial tropism in a *de novo* host-symbiont association by transinfecting the systemic host *D. simulans* (STC) with the *Wolbachia* strain *w*Wil from *D. willistoni*, a representative of the restriction type, we thereon named *w*Wil/STC (**Table 1**). For the sake of simplicity in the following text, we use SIT and RIT abbreviations to define <u>s</u>ystemic infection type and restricted infection type, respectively.

**Table 1.** *Drosophila* species and lines used in the study.

| *Drosophila* species | subgroup | line code | short name | *Wolbachia* strain |
| --- | --- | --- | --- | --- |
| *D. melanogaster* | melanogaster | Harwich H2 | MEL | wMel |
| *D. simulans* | melanogaster | KB30STC | STC | wAu |
| *D. tropicalis* | willistoni | Trop1 | TRO | wTro |
| *D. paulistorum* | willistoni | Pau5 O11 | PAU | wPau |
| *D. willistoni* | willistoni | JS6.3 | WIL | wWil |
| *D. septentrionalis* | saltans | SEP1/PLR | SPT | wSpt |
| *D. sturtevantii* | sturtevantii | FG707 | STV | wStv |
| *D. simulans* TI<sup>§</sup> | melanogaster | wilE/STC 36 | wilE/STC | wWil |

<sup>§</sup> transinfected by microinjection

### Tropism of Wolbachia in adult and larval nervous tissues of Drosophila

We conducted fluorescent *in situ* hybridization (FISH) analysis using *Wolbachia*-specific 16S rRNA probes to study the bacterial distribution in adult brains of all six species listed above. As shown in **Figure 1A-C**, *D. septentriosaltans* (SPT) and *D. tropicalis* (TRO) exhibit, similar to *D. melanogaster* (MEL), a SIT pattern with bacteria evenly distributed all over the tissue without accumulation in certain brain regions. In contrast, *Wolbachia* of *D*. *paulistorum* (PAU), *D. willistoni* (WIL) and *D. sturtevanti* (STV) were found to be locally restricted (**Figure 1D-F**). Although we did not focus on deciphering the identity of infected brain regions in the present study, all three species exhibited clear isolation of infection in certain regions of the brain, whereas most of the tissue was free of *Wolbachia*. For measuring *Wolbachia* tropism in respective brains, we determined the restriction indices (RI) as a number of uninfected cells divided by total number of cells (see Materials and Methods section). The indices revealed two significantly distinct groups of either systemic (MEL, SPT, TRO hosts) or restricted (PAU, WIL, STV hosts) infections (**Figure 1M**) with RI ranging from 0.02 to 0.12 and 0.82 to 0.88, respectively (Poisson regression: *p*<0.001).

**Figure 1.**
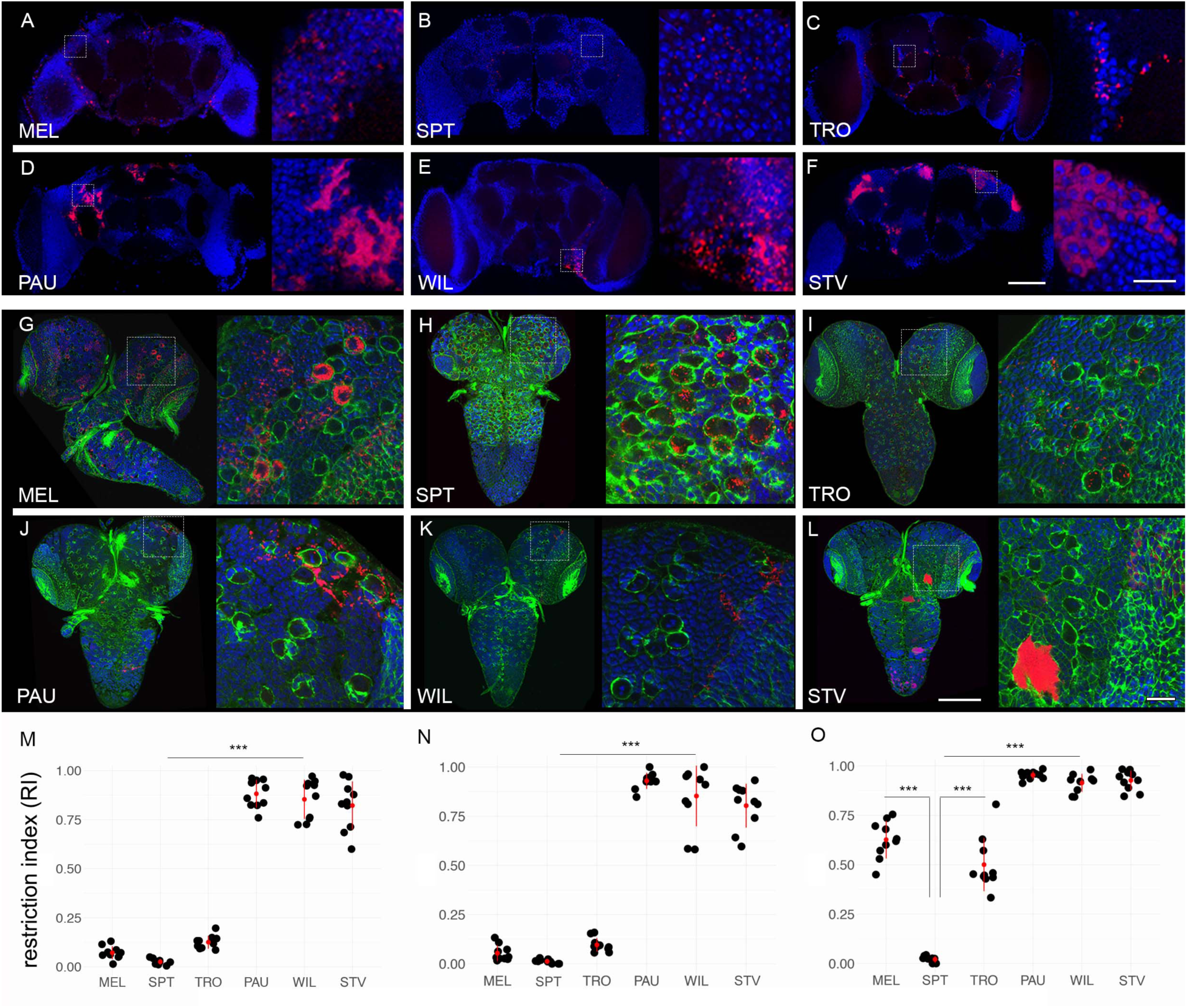
Restriction of *Wolbachia* infection in nervous tissues of neotropical *Drosophila*. Fluorescent *in situ* hybridization on different *Drosophila* adult brains (**A-F**) and 3^rd^ instar larval CNS (**G-L**) using 16S rRNA *Wolbachia*-specific probe (red). The bottom plots show restriction indices of all six species for *Wolbachia* infections in adult brains (**N**) and larval CNS (**M**), respectively. **O** shows RI of bacterial infection in neuroblasts of 3^rd^ instar larval CNS. DNA is stained with DAPI (blue) and actin with Phalloidin (green). For each *Drosophila* species 10 organs from each developmental stage were analyzed (Supplemental data file). Asterisks denote statistical significance (***, *p*<0.001; Poisson regression). Red bars show standard deviation, red dots designate the mean value. Scale bar: 50 μm.

Next, we examined the distribution of *Wolbachia* in the central nervous system (CNS) of 3^rd^ instar larvae. The analysis of bacterial infection in larvae of all six species (**Figure 1G-L**) using same FISH approach demonstrated similar results as obtained for the adult brains. The larval nervous tissue from MEL, SPT and TRO showed systemic infection (**Figure 1G-I**), whereas *Wolbachia* in PAU, WIL and STV were locally restricted (**Figure 1J-L**). Evaluation of the RI for *Wolbachia* infection revealed a limited restriction of bacteria in SIT species in which the index ranged from 0.01 to 0.09. Conversely, the high indices in RIT species ranged from 0.80 to 0.92 (**Figure 1N**; Poisson regression: *p*<0.001). Hence, the pattern of bacterial localization is already determined in the larvae and preserved through metamorphosis.

The nervous system of 3^rd^ instar larvae consists of three different cell types, i.e., neuroblasts (neural stem cells), neurons and glial cells (Homem and Knoblich, 2012). We therefore asked whether the endosymbiont targets any of these cell types specifically or acts regardless of the lineage in a locally restricted manner. Using a neuroblast-specific antibody against Deadpan, a transcriptional repressor responsible for maintenance of neuroblast’s self-renewing, and also a glia-specific antibody against Repo, a transcriptional factor expressed in glial cells, we analyzed the cell type specificity of *Wolbachia* localization in the CNS of larvae of all six lines (**Figure S1**).

We found infections of glial cells located in the cortex of the CNS in all six analyzed species. MEL, SPT and TRO showed systemic patterns, whereas bacteria in PAU, WIL and STV were locally restricted (**Figure S2**). The majority of bacteria, however, were concentrated in neuroblasts and neurons of the larval CNS. Neuroblasts, which we differentiated from other cell types by their bigger size of approximately 10 μm in diameter (see the insets of **Figure 1G-L**), showed distinctive *Wolbachia* infection patterns depending on the species analyzed (**Figure S3A**). Bacterial densities in a single neuroblast were quantified by dividing the bacterial load within the cell to the area of the cell’s cytoplasm (**Figure S3A**). The highest accumulation of bacteria in neural stem cells was observed in MEL and STV with both densities equating to 0.76. In contrast, TRO and SPT exhibited the lowest densities of 0.13 and 0.30, respectively. Unlike these species, the densities in neuroblasts of PAU and WIL showed an unusually high variance within individual larval CNS, ranging from either 0.2 to 0.79 (mean = 0.51) or 0.1 to 0.79 (mean = 0.57), respectively. High variance in these two restricting hosts might suggest that their respective *Wolbachia* strains only target a specific, yet undetermined subset of neuroblasts. Quantification of RI of bacteria in neuroblasts of all six semispecies (**Figure 1O**) revealed that despite the SIT patterns in MEL and TRO, approximately only half of their neural stem cells were infected with *Wolbachia*, whereas in SPT almost all neuroblasts were *Wolbachia*-positive (0.63, 0.51 and 0.02; Poisson regression: *p*<0.001). On the other hand, in all hosts with RIT patterns (PAU, WIL and STV) the RIs were significantly higher than in the systemic ones (0.95, 0.93 and 0.92; Poisson regression: *p*<0.001).

By using a specific antibody against Asense, a transcriptional factor expressed in type I but not type II neuroblasts, we further specified the cell type of infection (**Figure S4**). Type II neuroblasts divide symmetrically producing intermediate neural progenitors, which then divide asymmetrically to self-renew and generate a ganglion mother cell whereas type I neuroblasts divide asymmetrically and only once (Homem and Knoblich, 2012). As a result, type II neuroblasts generate a greater number of cells in the adult brain than type I. We hypothesized that infecting type II neuronal stem cells might be an opportunity for *Wolbachia* to achieve a broader spread. In all three species with SIT pattern, *Wolbachia* were found in both neuroblast types (**Figure S4**, first 3 rows). For hosts with RIT patterns, however, only type I neuroblasts were found infected with the endosymbiont (**Figure S4**, last 3 rows).

Furthermore, to analyze the aggregation of *Wolbachia* infection in the CNS, i.e., the formation of clusters of neighboring neurons bearing infections, we quantified the average number of infected neurons in groups (**Figure S3B**). Quantifications demonstrated the formation of big clusters of infected neurons in SPT, MEL and STV (21.1, 18.5 and 15.9 neurons on average per cluster, respectively) and smaller clusters in WIL, TRO and PAU (13.5, 9.5 and 7.2 neurons on average per cluster, respectively), without statistically significant differences between systemic and restring hosts (*p*>0.05).

In summary, we observe two distinct patterns of *Wolbachia* tropism in *Drosophila* nervous tissues, the systemic in MEL, SPT and TRO with an overall distribution of infection and the restricted in PAU, WIL and STV with isolation of infection to certain areas of the tissue. The pattern of infection is already determined in 3^rd^ instar larvae and preserved through metamorphosis with no tropism to a specific type of nerve cell, but detectable at higher densities in neuroblasts, the neural stem cells. In order to screen more saltans group representatives, *Wolbachia* FISH in neuronal tissues of *D. prosaltans* (saltans subgroup) and *D. lehrmanae* (sturtevanti subgroup) exhibited, similarly to SPT and STV hosts, either systemic or restricted patterns, respectively (**Figure S5**). Interestingly, bacterial densities within neural stem cells as well as their ability to aggregate vary among different *Drosophila* hosts irrespective to their diagnostic SIT and RIT patterns.

### *Tropism of* Wolbachia *in* Drosophila *ovaries*

For transovarial transmission, *Wolbachia* endosymbionts need to colonize the female germline. *Drosophila* ovaries consist of reproductive and somatic tissues. The nurse cells and the oocytes, originating from the germline stem cells, form the reproductive part. Conversely, the follicle cells, which ensheath the former, are derived from the somatic stem cell niche and represent the somatic part (Kirilly and Xie, 2007). Our systematic analysis of bacterial infections using FISH in the adult ovaries at stage 3-5 of all six species revealed that the majority of bacteria are associated with the reproductive part. However, they are also found in the soma but generally at lower levels (**Figure 2A-F**).

**Figure 2.**
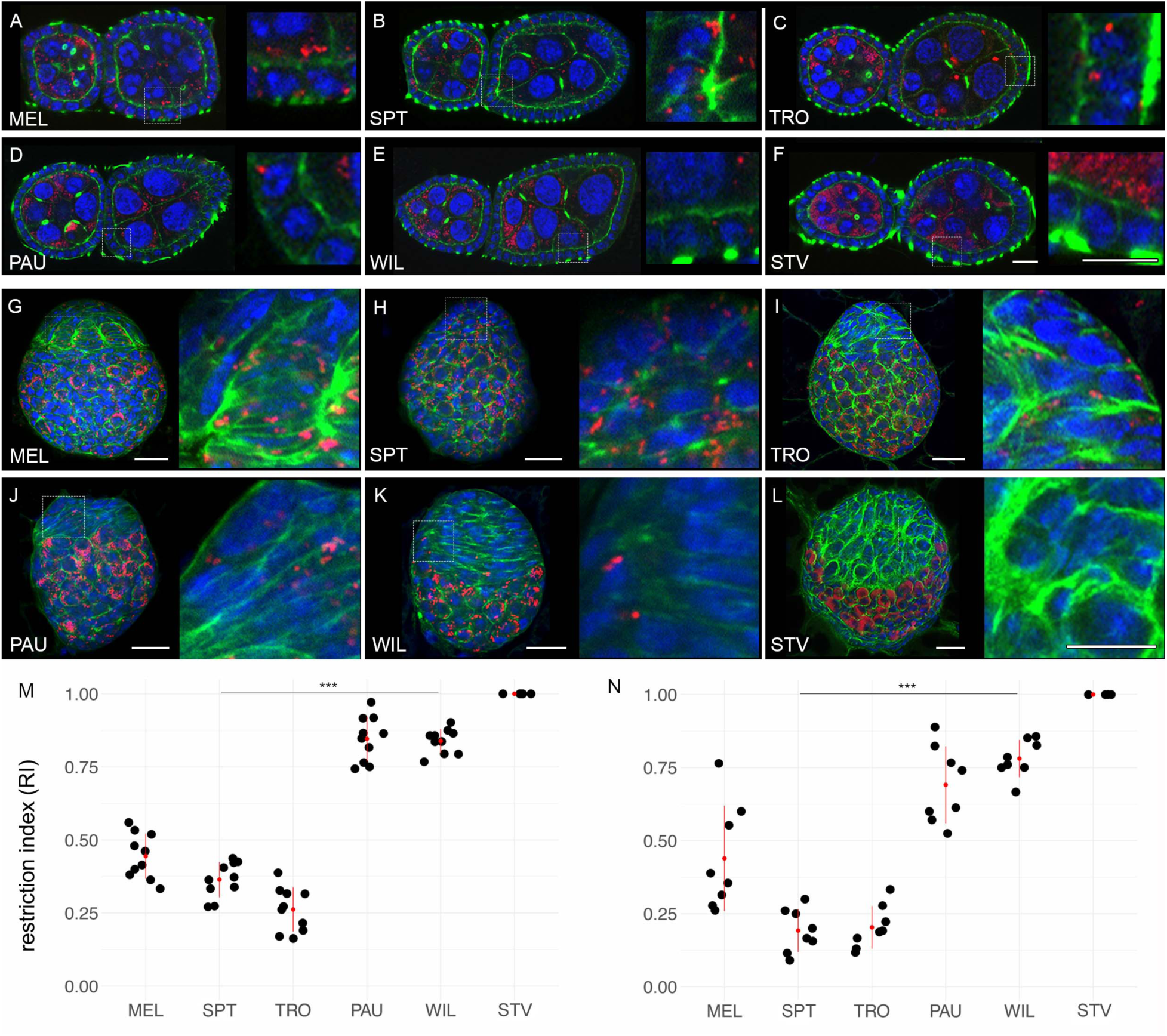
Restriction of *Wolbachia* infection in somatic and reproductive parts of adult and larval ovaries of neotropical *Drosophila*. Fluorescent *in situ* hybridization of different *Drosophila* adult ovaries (**A-F**) and 3^rd^ instar larval ovaries (**G-L**) using 16S rRNA *Wolbachia*-specific probe (red). RIs of *Wolbachia* infection in follicle cells of adult (**M**) and larval (**N**) ovaries for all six species. DNA is stained with DAPI (blue), actin with Phalloidin (green). Asterisks denote statistical significance (***, *p*<0.001; Poisson regression). Red bars show standard deviation, red dots designate the mean value. In total, 8-10 organs were analyzed for every species (Supplemental data file). Scale bar: 20 μm.

Infection density in the nurse cells and the oocyte of PAU, WIL and STV was significantly higher than in MEL, SPT and TRO (**Figure S6**; Poisson regression: *p*<0.001). We also observed *Wolbachia* infection in the somatic part of the ovaries. Respective RIs in follicle cells varied among the species with relatively low average values in the systemic hosts TRO, SPT and MEL (**Figure 2M**; 0.26, 0.36 and 0.44, respectively), but significantly higher in the restrictors WIL, PAU and STV (0.84, 0.85 and 1, respectively; Poisson regression: *p*<0.001).

The analysis of bacterial infection using FISH in 3^rd^ instar larval ovaries revealed similar results as observed in the adult ovaries (**Figure 2G-L**). The larval ovary can also be divided into somatic and reproductive parts either morphologically or by specific staining. Similar to adult ovaries, *Wolbachia* is dominating in the reproductive part (germ cells) of all six species analyzed. In the somatic part, low restriction of infection was observed only in systemic hosts SPT, TRO and MEL (**Figure 2N**; 0.19, 0.20 and 0.44, respectively), in contrast to significantly higher restriction in WIL and PAU (0.78 and 0.70, respectively; Poisson regression: *p*<0.001) and absence of infection in STV. The preservation of infection patterns in the somatic part of the adult ovary in comparison to the larval gonad is in accordance with the same trend described for the larval CNS and adult brain where the bacterial distribution was also preserved after metamorphosis.

### *Wolbachia* densities drop dramatically during early embryonic gastrulation in *Drosophila* species with restricting pattern of infection

Data obtained from the adult and 3^rd^ instar larval somatic and reproductive tissues demonstrate that cell type-specific tropisms of *Wolbachia* are determined already in larvae and are preserved during the metamorphosis of the host. To investigate how infection patterns form initially, we analyzed *Wolbachia* distribution through different *Drosophila* embryogenesis stages. Analysis of *Wolbachia* localization in early embryos (stage 3-5) revealed SIT patterns with no differences in infection distribution in any of the six tested hosts (**Figure 3**, left row). Bacteria were evenly dispersed all over the embryo and closely associated with the chromatin during mitosis. Interestingly, in mid-embryogenesis (stage 6-9), *Wolbachia* densities decreased in PAU, WIL and STV but not in MEL, SPT and TRO embryos (**Figure 3A**, middle row). Although bacteria were still evenly distributed across all embryonic areas in all six species at early gastrulation, many cells of PAU, WIL and STV embryo were already cleared of infection. Finally, at late embryogenesis (stage 13-15) we observed drastic differences in *Wolbachia* distribution between species with SIT and RIT patterns of bacterial infection (**Figure 3**, right row). While in systemic MEL, SPT and TRO hosts bacteria were equally dispersed in most embryonic tissues, *Wolbachia* in PAU, WIL and STV species were now locally restricted to the primordial germ cells (PGCs), the future gonads, and to some additional isolated somatic cell clusters in the embryo.

**Figure 3.**
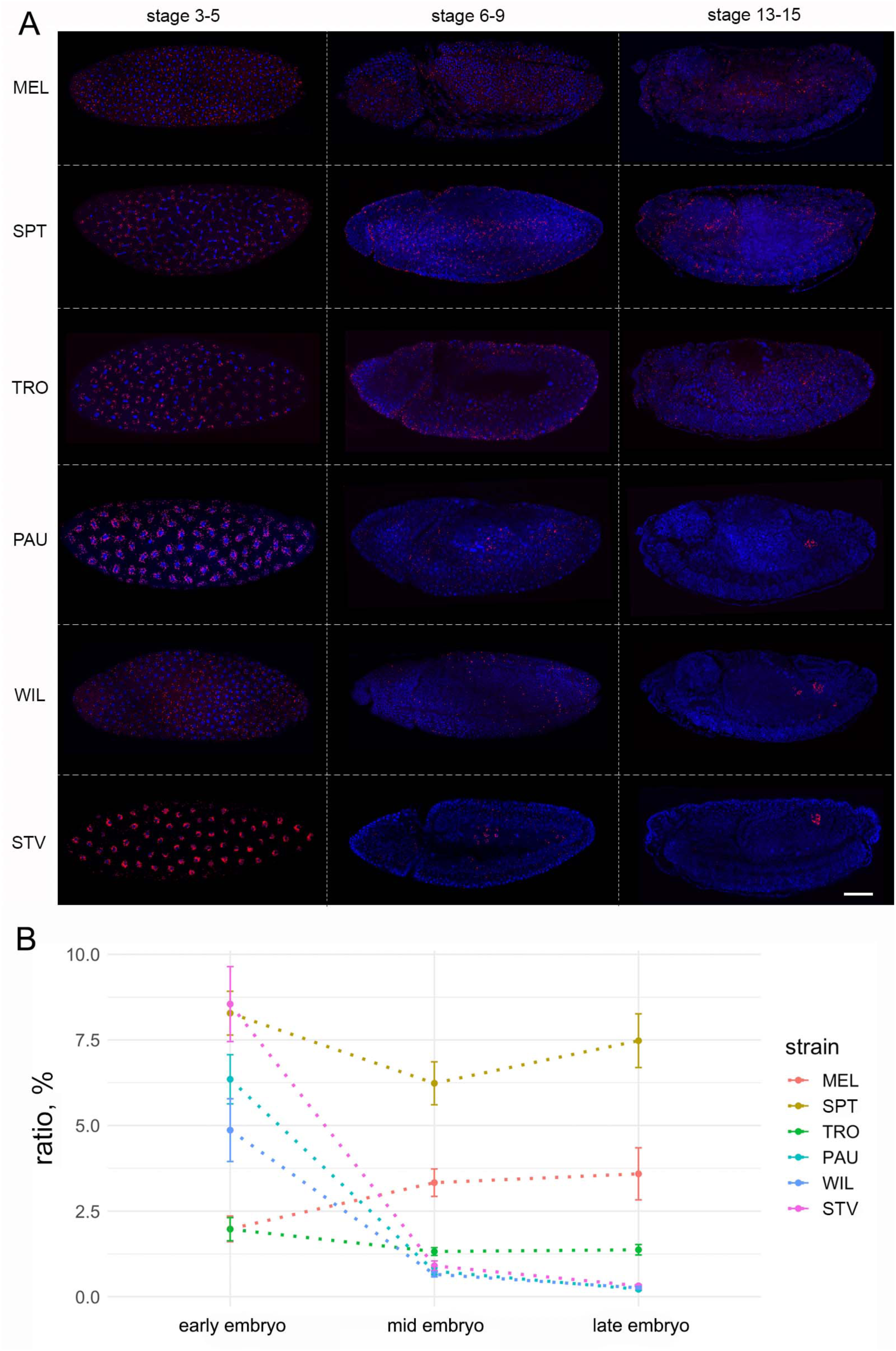
Dramatic reduction of *Wolbachia* density during mid-embryogenesis in neotropical *Drosophila* species. (**A**) Fluorescent *in situ* hybridization of *Drosophila* embryos at stage 3-5, 6-9 and 13-15 of embryogenesis, using 16S rRNA *Wolbachia*-specific probe (red). DNA is stained with DAPI (blue). (**B**) Quantification of *Wolbachia* density at early, mid and late embryogenesis, using Fiji, as bacterial density in a whole embryo divided by the area of an embryo. Bars show standard error of the mean. For each species and stage 5 embryos were analyzed for density measurements (Supplemental data file). Scale bar: 50 μm.

Quantification of global *Wolbachia* densities in embryos at these three defined developmental stages using Fiji confirmed this dramatic reduction of infection starting at mid-embryogenesis in PAU, WIL and STV (*p*<0.001, One-way ANOVA with Tukey HSD test), whereas densities of bacteria in MEL, TRO and SPT hosts remained unchanged across all stages (**Figure 3B**).

To verify our assumption that *Wolbachia* are selectively maintained mainly in PGCs of late WIL, PAU and STV embryos, we performed a sequential FISH and immunofluorescence analysis using an antibody against Vasa, a protein essential for the pole plasm assembly in the egg, which is commonly used as a germline precursor marker (Gustafson and Wessel, 2010). As expected from a maternally transmitted endosymbiont, all six tested host species harbored the bacterial infection within their PGCs (**Figure 4A**, left column). However, only PAU, WIL and STV hosts showed strict isolation of infection within the PGCs with infrequent bacterial localization in surrounding somatic tissue, whereas in MEL, SPT and TRO *Wolbachia* remained systemic (*p*<0.001; One-way ANOVA with Tukey HSD Test) (**Figure 4B**).

**Figure 4.**
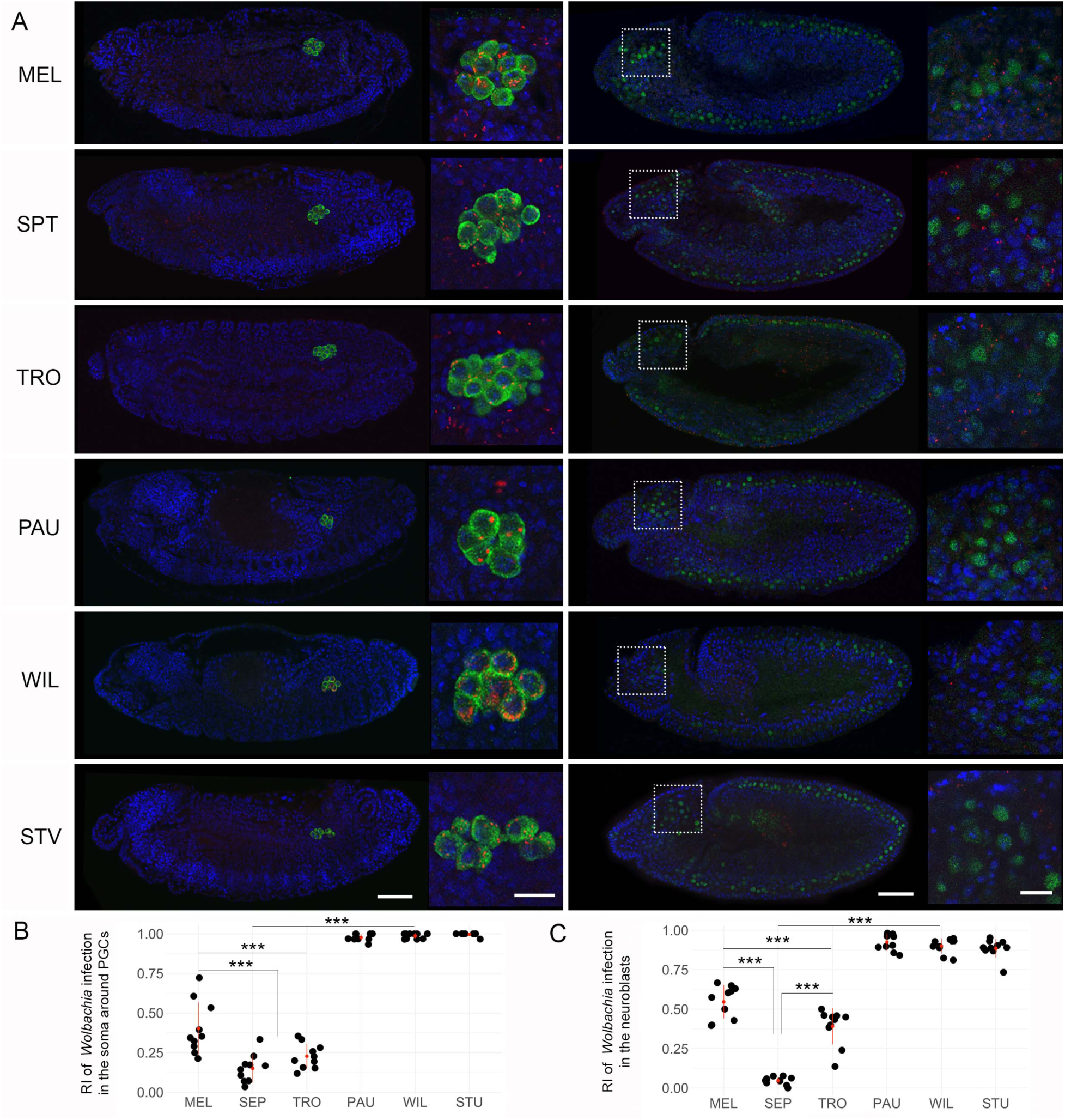
*Wolbachia* tropism to primordial germ cells and neuroblasts of neotropical *Drosophila* embryos. Sequential FISH using *Wolbachia*-specific 16S rRNA probe (red) and immunofluorescent staining of PGCs with anti-Vasa (left column, green) and neuroblasts with anti-Deadpan (right column, green) antibodies on *Drosophila* embryos. DNA is stained with DAPI (blue) (**A**). Determined RIs in the soma of neighboring PGCs (**B**) and in neuroblasts (**C**). In total ten embryos were analyzed for every cell type (Supplemental data file). Asterisks denote statistical significance (***, *p*<0.001; One-way ANOVA with Tukey HSD Test). Red bars show standard deviation, red dots designate the mean value. Scale bar: 50 μm for embryos, 10 μm for insets.

Additionally, using a similar approach but with the neuroblast-specific Deadpan antibody, we analyzed bacterial tropism in embryonic neuroblasts after their delamination from the neuroectoderm at stage 9-10 (**Figure 4A**, right column). Similar profound elimination of bacteria from somatic parts of the embryo (neuroblasts in this case) was observed in PAU, WIL and STV species in contrast to an ongoing systemic infection in MEL, SPT and TRO (*p*<0.001; One-way ANOVA with Tukey HSD Test). Already after delamination of the neuroblasts in pro-cephalic neurogenic region, which gives rise to the brain of an embryo, we detected only a very few nuclei associated with *Wolbachia* signals in species restricting the infection, whereas in the SIT hosts at least half of the neuroblasts contained the bacteria (**Figure 4A**, right column insets; **Figure 4C**).

In summary, by systematically tracing the temporal and spatial dynamics of *Wolbachia* tropism *in situ*, we found that bacterial densities started to drop already before gastrulation (stage 6-9) exclusively in three RIT species. The majority of *Wolbachia* accumulated mainly in PGCs but also in a few other cells of the embryo (neuroblasts and other undefined cell types). Hence, the restricted *Wolbachia* tropism found in the germline and the soma of PAU, WIL and STV flies is already determined before the onset of gastrulation, either by active host-directed elimination, or by dilution followed by selective replication of the native endosymbiont in some defined stem cells.

### Autophagy eliminates *Wolbachia* in restricting species during early gastrulation

Since we detected a dramatic decrease in bacterial titer already during embryogenesis, we hypothesized that active host-directed elimination of the endosymbiont is a more plausible mechanism of infection restriction than dilution and selective replication (**Figure 3B**). Autophagy was considered a potential mechanism for bacterial clearance because it has previously been demonstrated as a key cellular strategy for controlling *Wolbachia* density and tropism in *Brugia malayi* nematodes and *D. melanogaster* flies in vivo, as well as *in vitro* in cell lines of *D. melanogaster* and *Aedes albopictus* (Voronin et al., 2012). Moreover, it was recently shown that the density of *Wolbachia* in *D. melanogaster* is mediated by host autophagy in a cell type-dependent manner (Deehan et al., 2021). To test our hypothesis, we conducted sequential FISH and immunofluorescent analysis using an anti-GABARAP antibody, which is diagnostic for maturing autophagosomes in a cell. Since the drastic loss of somatic *Wolbachia* was clearly evident at mid-embryogenesis of restricted hosts (stage 6-9, see **Figure 3** middle row), we focused our analysis on early to late blastodermal embryos to study the temporal and spatial dynamics of the elimination process *in situ*. No signs of bacterial autophagy were found in somatic cells or in PGCs of systemic MEL, SPT and TRO hosts (**Figure 5 A-C**; **Figure S7**). However, in the soma of the restricted PAU, WIL and STV embryos, we clearly observed the formation of GABARAP-positive rings around bacterial cells (**Figure 5D-F**). The earliest cases of *Wolbachia* engulfment were detected in blastodermal embryos (stage 5), with the highest peak in early gastrulation (stage 6) and only rarely at later stages (stage 7-8). Importantly, PGCs, which could be clearly recognized as an isolated cell cluster at posterior part of the embryo in late blastodermal or early gastrulating embryo, were devoid of any signs of bacterial autophagy in all three species with the restricted pattern (**Figure 5G-I**). This was in full agreement with our observations in later embryos that *Wolbachia* is preserved and maintained in the gonad precursor cells (**Figure 4A**, left column).

**Figure 5.**
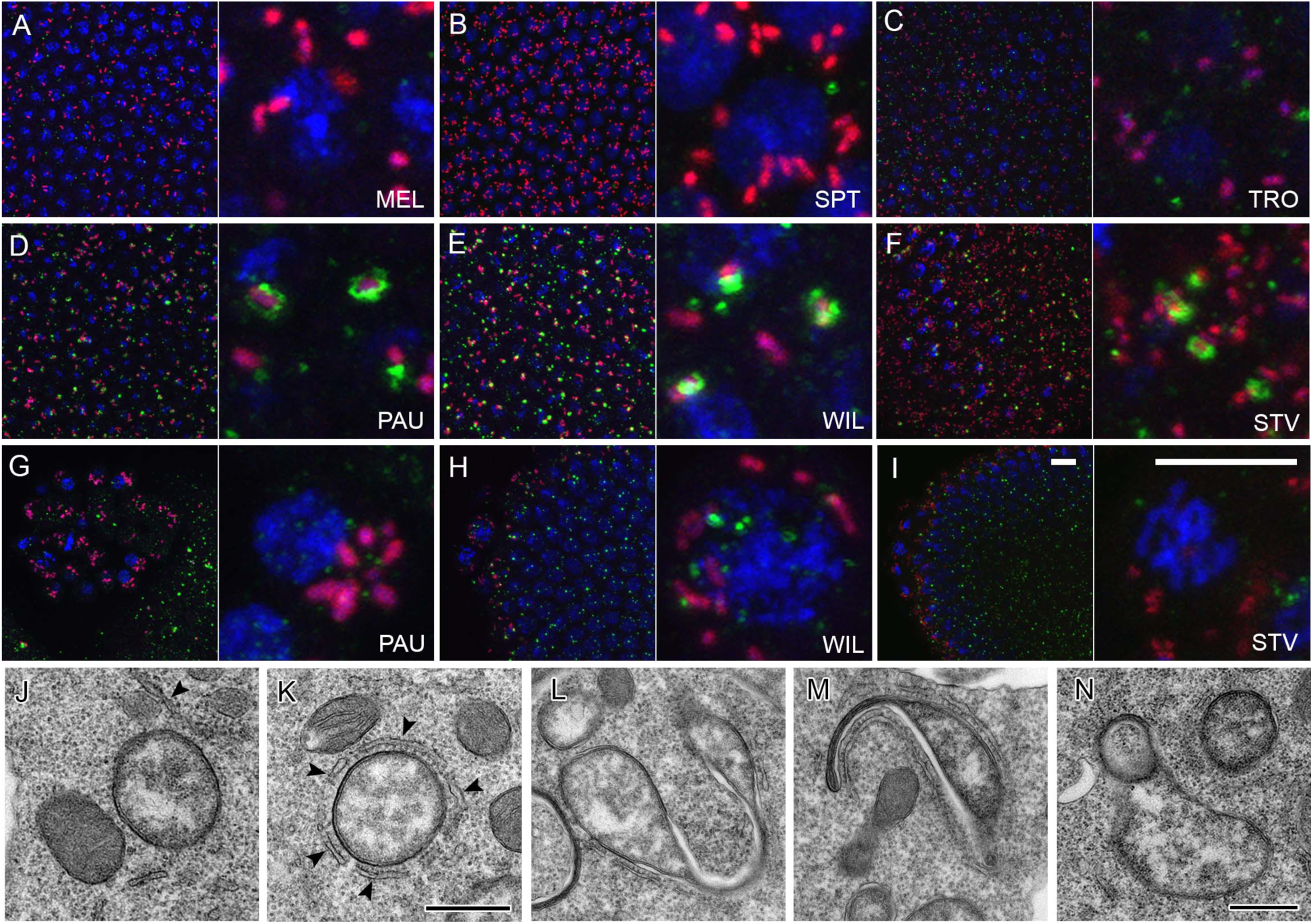
Elimination of *Wolbachia* via autophagy in neotropical *Drosophila* embryos. Sequential FISH using *Wolbachia*-specific 16S rRNA probe (red) and immunofluorescent staining with anti-GABARAP (green) antibody of embryos at stage 5 (**A-I**). Note the absence of autophagy in SIT species (**A-C**) and formation of autophagosomes (green rings) around *Wolbachia* in RIT species (**D-F**). Also note the absence of autophagy in PGCs of RIT species (**G-I**). Transmission electron microscopy on systemic MEL (**J**) and restrictive PAU (**K**) embryos at the cellularization and early gastrulation (stage 5-6). Contrary to MEL (**J**), tight physical associations between *w*Pau *Wolbachia* and the endoplasmic reticulum of restrictive PAU hosts (arrowheads) are prominent (**K**). Abnormal *w*Pau *Wolbachia* morphotypes (**L-N**) with signs of stretching (**L**), membrane extrusions (**M**) and vesicle formation (**N**). DNA is stained with DAPI (blue). Scale bar: 10 μm for all fluorescent images, 0.5 μm for TEM.

To further support our observation, we quantified the co-localization of GABARAP and FK2 antibodies and *Wolbachia* cells using a JACoP plugin (Bolte and Cordelieres, 2006) for the imaging software Fiji (Shindelin et al., 2012). We found a pronounced overlap of autophagosomes and *Wolbachia* in somatic parts of the blastodermal and early gastrulating embryos (stage 5-6) of PAU, WIL and STV species with 22.3 ± 2.2%, 25.8 ± 3.4% and 15.5 ± 4.1%, respectively. By contrast, in somatic parts of earlier embryos (stage 3-4) and PGCs at both developmental stages of all six species we detected significantly less co-localization (between 0 and 2%) of *Wolbachia* with the antibody (Poisson regression: *p* < 0.001), confirming that there is no clearance of bacterial infection at this stage (**Figure S8A**).

To further decipher the mechanistic basis of these intimate bacterial interactions with autophagosomes, we conducted an ultrastructural analysis of MEL and PAU embryos at cellularization and early gastrulation stage. Transmission electron microscopy (TEM) of PAU embryos at these stages revealed intimate interaction of *Wolbachia* with the endoplasmic reticulum (ER) of the host cell, contrary to MEL species, where no similar types of tight associations were detected (**Figure 5J, K**). In most of the cases we observed rough ER membranes encircling the bacterial cells by close apposition but without direct contact (**Figure 5K**, **Figure S9**). Later in early gastrulating PAU embryos, abnormal *Wolbachia* bacteria are dominant, exhibiting various signs of stretching, membrane extrusions and vesicle formation (**Figure 5L-N**, **Figure S9A-C**) that indicate symbiont degradation. No such structures were observed in MEL embryos at this stage. Surprisingly, we did not observe any autophagosome-like structures or traces of lysed bacteria at cellularization and early gastrulation, which is in contrast to clear co-localization of anti-GABARAP antibody and *Wolbachia* obtained with sequential FISH and immunofluorescent staining (**Figure 5D**, **Figure S8A**). The most plausible explanation of this observation is that autophagy of bacteria occurs in a non-canonical way. The abnormal *Wolbachia* forms we detected in early gastrulating embryos of restricting species support this hypothesis.

Besides anti-GABARAP, we also used an anti-FK2 antibody, which recognizes mono- and poly-ubiquitinated conjugates, to decipher whether bacteria are tagged for subsequent degradation. Consistent with our previous observations with anti-GABARAP staining, we did not detect any signs of ubiquitination of *Wolbachia* in MEL, SPT and TRO embryos at blastodermal and gastrulating stages (**Figure S9D-F**), including the PGCs (**Figure S7**). Furthermore, we did not detect frequent co-localization of anti-FK2 antibody and *Wolbachia* in PAU and STV embryos at both embryonic stages (**Figure S9G, I** and **Figure S8B**). Surprisingly, only WIL embryos exhibited pronounced ubiquitination signals associated with *Wolbachia* already at the blastodermal stage of embryogenesis (**Figure S9H**). The signal from the antibody staining was confined on one halve of the bacterial surface, in contrast to the “ring”-like structures observed with anti-GABARAP (**Figure 5E**).

To sum up, analysis of blastodermal and early gastrulating embryos revealed that *Wolbachia* are most likely eliminated from the tissues of restricting hosts by autophagy mediated by intimate interactions of ER membranes with bacterial cells. While *w*Wil bacteria are tagged and presumably degraded in a ubiquitin-dependent manner, the two other native endosymbionts of PAU and STV are eliminated in a slightly different and most likely ubiquitin-independent way. The mechanistic basis of these observed differences awaits further studies in our laboratory.

### Host background plays a major role in regulating the pattern of *Wolbachia* tropism in the soma

To test the influence of each partner in this intimate symbiotic association, we conducted experiments with transinfected flies carrying different *Wolbachia* strains in the same host background. *Drosophila simulans* flies that are naturally infected with *Wolbachia* strains like *w*Au or *w*Ri, demonstrating the SIT, were first cleared from the infection using antibiotics (now named *D. simulans* STC), and subsequently transinfected with *w*Wil strain from *D. willistoni* via embryonic microinjections. Thus, a *Wolbachia* strain accommodated to the restricting host background was introduced into the SIT environment. In our experiment, the successfully transinfected line *w*Wil/STC was kept in the lab for more than 10 years before starting further analyses on symbiont tropism in the *de novo* host background. Comparative FISH analysis of 3^rd^ instar larval CNS and adult ovaries (stage 3-5) with *Wolbachia*-specific probes showed that the *de novo w*Wil infection in *D. simulans* is not restricted as in *D. willistoni*, but systemic, similar to the globally dispersed patterns when infected with their natural strains of *Wolbachia* (**Figure 6A**).

**Figure 6.**
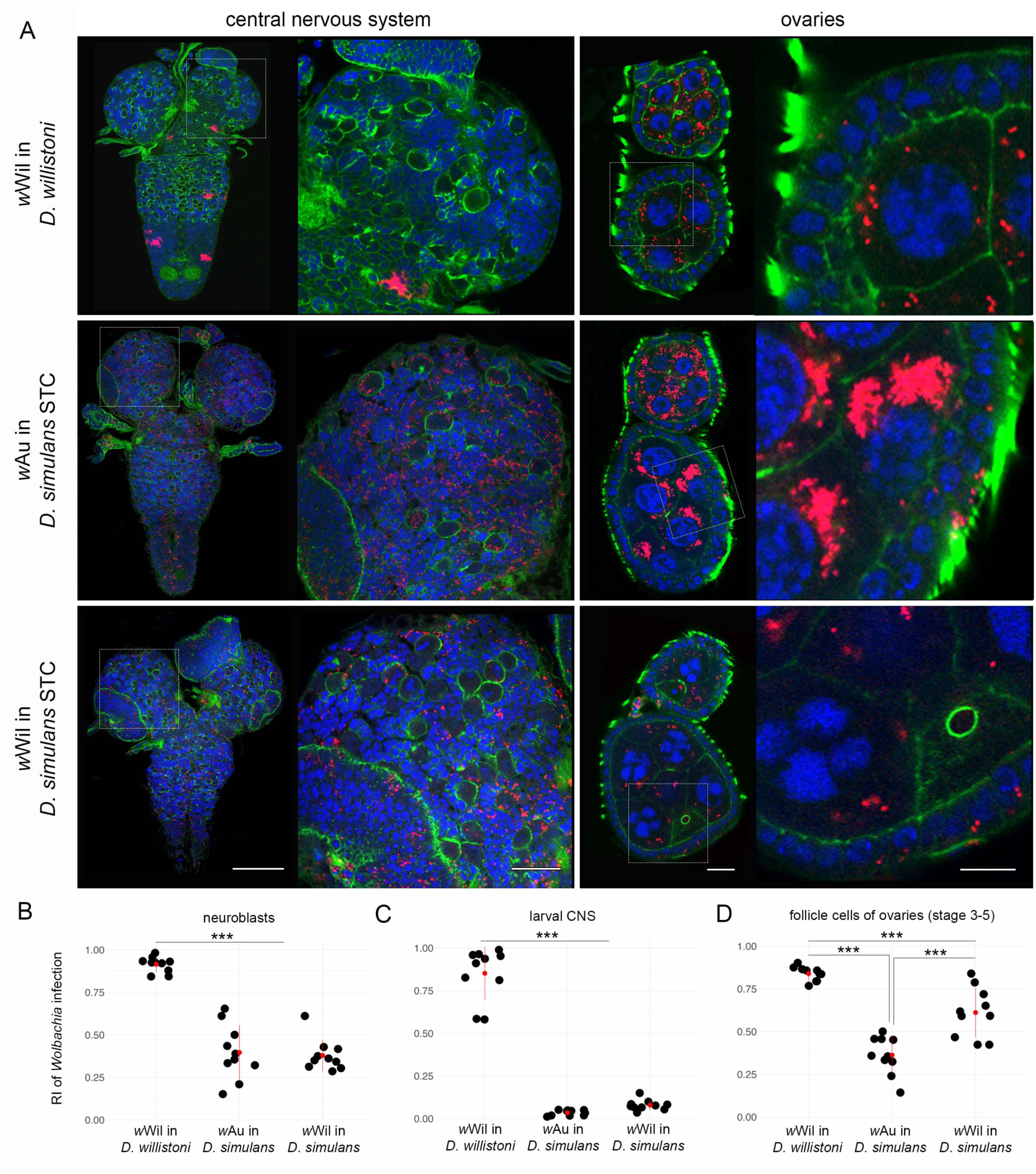
Tropism of the restrictive *w*Wil strain from *D. willistoni* in systemic *D. simulans* host. Fluorescent *in situ* hybridization of different *Drosophila* 3^rd^ instar larval CNS (**A**, left column) and adult ovaries at stage 3-5 (**A**, right column) of *D. willsitoni*, *D. simulans* and *D. simulans* transinfected with *w*Wil strain using 16S rRNA *Wolbachia*-specific probe (red). **B** demonstrates the RI of bacteria in neuroblasts. **C** and **D** show the RIs of *Wolbachia* infection in the larval CNS and follicle cells of adult ovaries, respectively. DNA is stained with DAPI (blue); actin is stained with Phalloidin (green). For each *Drosophila* species 10 organs from each developmental stage were analyzed (Supplemental data file). Asterisks denote statistical significance (***, *p*<0.001; One-way ANOVA with Tukey HSD Test). Red bars show standard deviation, red dots designate the mean value. Scale bar: 20 μm.

Quantification of the RI for infection of neuroblasts and whole larval CNS in *w*Wil/STC (**Figure 6B, C**) confirmed the systemic nature of *w*Wil localization in *D. simulans* with no difference to native *w*Au in *D. simulans* (*p*=0.93 for neuroblasts and *p*=0.52 for larval brains, One-way ANOVA with Tukey HSD Test), contrary to highly restricted tropism of *w*Wil in its native *D. willistoni* background (*p* < 0.001, One-way ANOVA with Tukey HSD Test). Interestingly, the infection of follicle cells in the adult ovaries of transinfected *w*Wil/STC flies was found to have a medium RI (**Figure 6D**) compared to systemic *w*Au in *D. simulans* (*p*<0.001, One-way ANOVA with Tukey HSD Test) and the highly restricted *w*Wil strain in *D. willistoni* (*p* <0.001, One-way ANOVA with Tukey HSD Test). Sequential FISH with *Wolbachia*-specific probes and immunofluorescence using anti-GABARAP and anti-FK2 antibodies on early embryos showed in contrary to *w*Wil in *D. willistoni* no physical interaction of native *w*Au and *de novo w*Wil with autophagosomes and the absence of ubiquitination in *D. simulans* hosts (**Figure 7A, B**, respectively). This observation was confirmed by quantitative co-localization of *Wolbachia* and the antibody signal using JACoP plugin in Fiji (**Figure S10)**.

**Figure 7.**
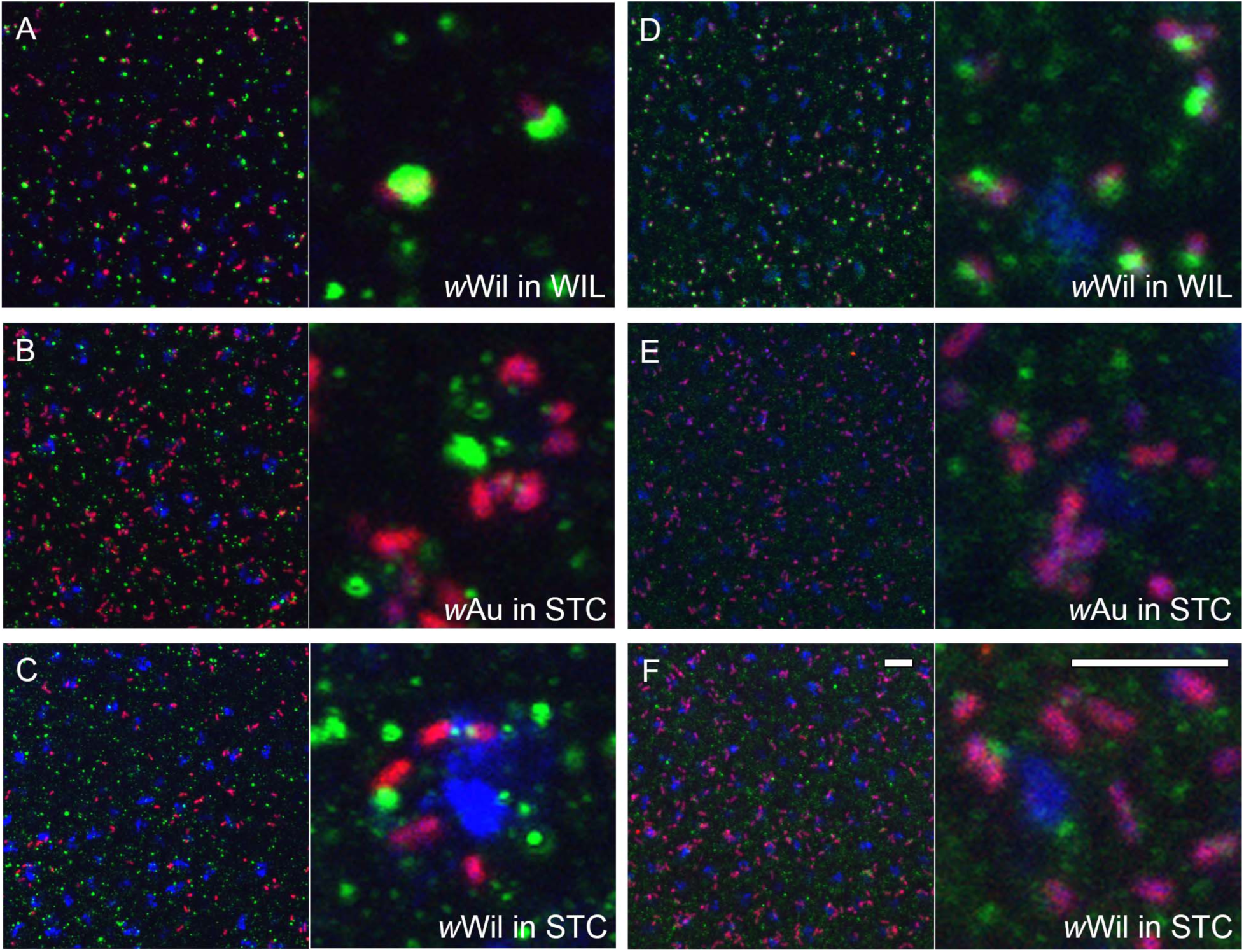
*Wolbachia* interactions with the host cell. Sequential FISH using *Wolbachia*-specific 16S rRNA probe (red) and immunofluorescent staining with anti-GABARAP (**A-C**) and anti-FK2 (**D-F**) antibodies on stage 6 embryos from *D. willsitoni* (*w*Wil in WIL), natively *w*Au-infected *D. simulans* (*w*Au in STC) and *w*Wil-transinfected *D. simulans* (*w*Wil in STC) lines. DNA is stained with DAPI (blue). Scale bar: 10 μm.

In summary, we observe that the host background plays a major role in regulating the distribution of the endosymbiont in its tissues.

## Discussion

In our previous study (Strunov et al., 2017), we demonstrated the remarkable restriction of *Wolbachia* bacteria to certain areas of the adult and larval central nervous system of *D. paulistorum* flies, which is in stark contrast to *D. melanogaster* and other insect hosts that usually harbor systemic bacterial infections in neuronal tissues (Min and Benzer 1997; Albertson et al., 2013; Strunov and Kiseleva, 2016). We hypothesized that the restricted tropism plus laterality of the endosymbiont to defined *D. paulistorum* brain regions might have evolved in order to keep the obligate mutualistic *Wolbachia* – *D. paulistorum* symbiosis in balance in a cost-benefit equilibrium, since it is essential for host’s oogenesis and directs mating behavior of both sexes (Miller et al., 2010; Schneider et al., 2019).

To survey *Wolbachia* infection patterns more broadly, we analyzed bacteria-host interactions with focus on tropism by comparative and quantitative FISH analyses in several additional neotropical *Drosophila* species belonging to the willistoni and saltans species group. Based on sequence similarities, *Wolbachia* in both groups appear to be closely related to *w*Au- like strains, with the exception of *w*Stv of *D. sturtevanti*, which differs significantly from the others (Bayraktar et al., 2010; Riegler et al., 2012; Martinez et al., 2021). In our present study we found that similar to *w*Pau in *D. paulistorum*, native *w*Wil *Wolbachia* are locally restricted in larval and adult brains, whereas *D. tropicalis*, a close relative to *D. willistoni*, exhibits clear patterns of the SIT, similar to *w*Mel in *D. melanogaster*. In *D. septentriosaltans*, a representative of the saltans species group, we found no signs of tropism in host flies carrying the *w*Spt *Wolbachia* strain that also belongs to the *w*Au-like group (Miller and Riegler, 2006; Riegler et al., 2012). In *D. sturtevanti*, however, *w*Stv *Wolbachia* are locally restricted similarly to the RIT of *w*Pau and *w*Wil in native willistoni group hosts. Interestingly, the characteristic restriction pattern of *w*Stv is also conserved in the closely related and newly described species *D. lehrmanae* (Madi-Ravazzi et al., unpublished) that carries a similar *w*Stv-like *Wolbachia* strain (Miller, unpublished).

### Tissue-tropism of Wolbachia has evolved at least twice in neotropical Drosophila hosts

In the current study we uncovered RIT patterns of the endosymbiont in three neotropical *Drosophila* hosts belonging to two different species groups that carry either *w*Au- or *w*Stv-like *Wolbachia* variants. This finding suggests that the local restriction of the endosymbiont evolved at least two times independently in neotropical *Drosophila* by targeting two different *Wolbachia* variants – the closely related and more ancestral *w*Au-like strain in the lineage of *D. paulistorum* and *D. willistoni*, and the more recently acquired *w*Stv-like bacteria of *D. sturtevanti* and *D. lehrmanae*. As *w*Au-like *Wolbachia* are conspecific and the dominating, most likely ancestral, infection type of neotropical *Drosophila* species (Miller and Riegler, 2006) we speculate that the last common ancestor of *D. sturtevanti* and *D. lehrmanae* might have carried a *w*Au-like strain too, which in the following got lost in competition with the arrival and successful establishment of the newly acquired *w*Stv stain. Under the assumption that the ancestral *w*Au infection was similarly restricted to defined tissues as *w*Wil and *w*Pau in their native willistoni group prior to *Wolbachia* strain replacement, we hypothesize that the newly arrived and possibly more aggressive *w*Stv variant became domesticated and attenuated in the same way as the ancestral *w*Au-like infection type before. By this, the host was already pre-adapted to costly *Wolbachia* infections by restricting and limiting the endosymbiont to defined somatic niches where the cost-benefit equilibrium was not disturbed. In order to test this hypothesis, however, more data on *Wolbachia* tropism will be essential from more species of the saltans group since to date only systemic infections of *w*Au-like strains were found in *D. septentriosaltans* (this study) and *D. prosaltans* (Strunov, unpublished).

### Wolbachia tropism in adults is already determined in early embryos

Our comparative studies performed by systematic *Wolbachia*-specific FISH uncovered that adult *D. paulistorum* and *D. willistoni* as well as *D. sturtevanti* flies, all natively infected by either *w*Au- or *w*Stv-like strains, share similar patterns of local symbiont restrictions in their respective brains and ovaries. This RIT tropism is already manifested in early-mid embryogenesis by local restriction of the endosymbiont to the PGCs of the future germline and a few cell clusters of the soma (including neuroblasts) suggesting that both stem cell types might serve as the infection reservoir for the future imago.

We hypothesize that the massive reduction of bacterial titer in early embryogenesis is necessary to alleviate the burden of infection for the adult fly establishing the cost-benefit equilibrium in the system, since systemically infected species of PAU, WIL and STV were never observed in the lab as well as in recently collected wild specimens from French Guiana (data not shown). Semi-quantitative analyses of bacterial densities during early embryogenesis demonstrated that all three neotropical *Drosophila* with RIT patterns exhibit high titer *Wolbachia* infections (**Table 2**). In *D. tropicalis*, a close relative of *D. paulistorum* but exhibiting the SIT, *Wolbachia* titer in early embryos is stably low, and only slightly higher than in PAU, WIL and STV at later stages after bacterial elimination.

**Table 2.** Summarized characteristics of *Wolbachia* strains in native and novel hosts analyzed in the present study.

| *Wolbachia* strain/*Drosophila* species | *Wolbachia* distribution pattern | *Wolbachia* titer in embryogenesis |  |  | *Wolbachia* titer in the nurse cells and oocyte | *Wolbachia* titer in the neuroblasts | The number of infected cells in late embryogenesis | *Wolbachia* transmission efficiency to the daughter cell |
| --- | --- | --- | --- | --- | --- | --- | --- | --- |
|  |  | early | mid | late |  |  |  |  |
| wMel/*D. melanogaster* | systemic | ++ | ++ | ++ | + | +++ | +++ | ++ |
| wSpt/*D. septentrionalis* | systemic | +++ | +++ | +++ | + | + | +++ | +++ |
| wTro/*D. tropicalis* | systemic | ++ | ++ | ++ | + | + | +++ | ++ |
| wPau/*D. paulistorum* | restricted | +++ | ++ | + | +++ | ++ | + | + |
| wWil/*D. willistoni* | restricted | +++ | ++ | + | +++ | ++ | + | + |
| wStv/*D. sturtevantii* | restricted | +++ | ++ | + | +++ | +++ | + | + |
| wAu/*D. simulans* | systemic | ++ | ++ | ++ | +++ | + | +++ | ++ |
| wWil/*D. simulans* | systemic | ++ | ++ | ++ | ++ | + | +++ | ++ |

*Wolbachia* densities in embryos are strain-specific and being set in the unfertilized eggs during oogenesis (Serbus et al., 2008). After fertilization during the early nuclear divisions, they presumably do not replicate but only segregate (Lassy and Karr 1996; Miller unpublished). Thus, it seems likely that the smaller numbers of *Wolbachia* observed in early staged embryos of *D. tropicalis* are possibly below a critical threshold and less costly in hosts with SIT. In RIT hosts, higher densities seem detrimental and are hence avoided by elimination from most somatic parts of the embryo, which by natural selection leads to endosymbiont’s restriction in the host. In contrast to *D. tropicalis*, in *D. septentriosaltans*, another species with systemic *Wolbachia* infection, the bacterial titer is stably high in embryogenesis, however, at later developmental stages and especially in the imago the infection density decreases to MEL and TRO levels (**Table 2**). This reduction might occur due to a dilution effect via endosymbiont dissemination all over the developing organism during multiple cell divisions. In line with this idea, we previously demonstrated that some *D. paulistorum* semispecies harbor so-called low-titer *Wolbachia* infections (Miller et al., 2010) that are under the detection limit of standard PCR methods and hence more sensitive methods are needed for their identification (Arthofer et al., 2009; Miller et al., 2010; Schneider et al., 2014; Schneider et al., 2019; Baião et al., 2019).

We propose two main criteria for the establishment of *Wolbachia* tropism in symbiotic association: (i) the number of infected cells in late embryogenesis as a foundation of infection and the efficiency of *Wolbachia* transmission into dividing daughter cells during mitosis (**Table 2**). The first criterion represents a starting point with determined bacterial densities and localization, which is set in early-mid embryogenesis. In RIT hosts it occurs via directed elimination of bacteria from most somatic parts of the embryo and each infected pluripotent stem cell like PGC or neuroblast can be considered as a niche for the endosymbiont. The second criterion determines the future pattern of *Wolbachia* tropism in the adult fly by dissemination of infection from the niches by mitosis during development. The data on *Wolbachia* distribution in the nervous tissue of different *Drosophila* species across development demonstrated in this study and previously published (Albertson et al., 2009; Strunov et al., 2013) support this idea (summarized in **Figure S11**). In RIT hosts, the number of infected embryonic neuroblasts in the delaminated neuroectoderm is low due to extensive overall elimination of *Wolbachia* in the soma earlier in embryogenesis (**Figure S11A-C**). Later in development, these restricted infection niches give rise to clusters of bacterial infection in the larval CNS and adult brains, which differ in sizes depending on the transmission efficiency (**Figure S11A-C**). In the two systemic species with SIT, MEL and TRO, the ratio of infected neuroblasts is around 50% but the transmission efficiency is high enough to form multiple clusters of infection generating the SIT pattern (**Figure S11D, E**, respectively). In some species, not described in the present study, the dissemination of infection from the niches might be close to zero thus occupying only neuroblasts (**Figure S11F, I**). Finally, in SPT flies that also exhibit SIT the number of infected neuroblasts is almost 100% and the efficiency of transmission is high, which leads to overall dissemination of infection in the adult fly (**Figure S11G, H**).

The *Wolbachia* transinfection experiment, bringing *w*Wil bacteria from the RIT host *D. willistoni* into the SIT background of *D. simulans*, demonstrated that mainly the host background regulates the distribution pattern of infection in somatic tissues. These data are not entirely consistent with previous results for different *Drosophila* tissues, where in most cases the *Wolbachia* strain determined the tropism (summarized in **Table S1**). Such a discrepancy might be explained by different *Wolbachia* strategies to infect reproductive and somatic tissues. For instance, our data demonstrated that *Wolbachia* localization pattern is not strictly regulated by the host in follicle cells of adult ovaries from transinfected line (*w*Wil/STC).

### Autophagy is a key mechanism, eliminating Wolbachia during early Drosophila embryogenesis

Understanding the host-symbiont interaction regarding tropism and density control in the *Wolbachia*-*Drosophila* model system is of great importance for deciphering the essence of inter-kingdom relationships, which could be also applied to *Wolbachia*-mosquito and other symbiotic associations.

In three out of six *Drosophila* species analyzed in the present study we observe high restriction of *Wolbachia* to certain areas in some somatic tissues and their accumulation in reproductive organs of the host. This restriction occurs in early embryogenesis during the narrow time window between cellularization (stage 5) and early gastrulation (stage 6-7) with the infection being substantially reduced in the somatic part but staying high in PGCs. This massive somatic elimination of *Wolbachia* coincides with maternal-to-zygotic transition in *Drosophila* embryogenesis, which is marked by extensive degradation of deposited maternal mRNA and activation of zygotic gene expression (Tadros and Lipshitz, 2009). In this study we were able to dissect the process of *Wolbachia* clearance stepwise and demonstrate that bacteria are removed from the soma of RIT embryos via autophagy, which is schematically summarized in **Figure 8**. To our knowledge, this is the first example of autophagy-mediated regulation of bacterial densities during early development of the host.

**Figure 8.**
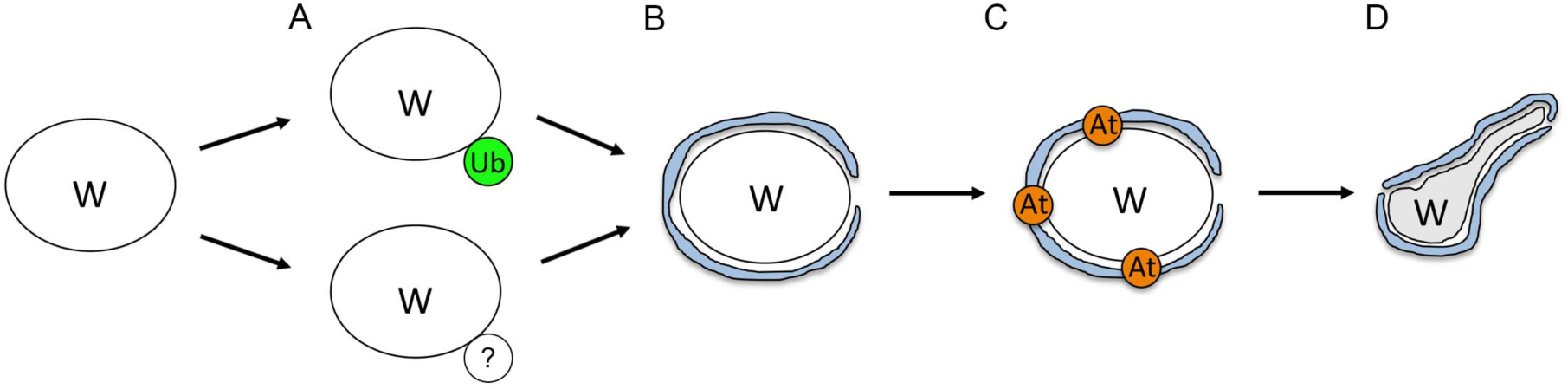
A scheme of *Wolbachia* elimination process during early host embryogenesis. **A** denotes the first step in infection elimination - ubiquitination (Ub), which is active in WIL hosts and absent in PAU and WIL. **B** demonstrates a second step - the encircling of the bacteria by ER membranes. **C** shows a third step – the attraction of autophagy machinery to the vesicle formed by ER. **D** depicts the last step – degradation of bacteria through yet undescribed mechanism

We propose that the first step of the bacterial elimination process is ubiquitination of the endosymbiont (**Figure 8A**). It is generally used by cells to tag proteins for proteasomal degradation (Weissman, 2001), but is also known for targeting intracellular bacteria for further elimination via autophagy during cellular defence against infections (Fujita and Yoshimori, 2011). In our study, however, we observe co-localization of ubiquitin with *Wolbachia* only in WIL species, whereas the other two RIT hosts PAU and STV showed low or no signs of it. Near absence of co-localization of ubiquitin with the native endosymbionts suggests that in these two hosts *Wolbachia* elimination occurs through ubiquitin-independent pathway (Khaminets et al., 2016). In contrast to *w*Wil, *w*Pau and *w*Stv *Wolbachia*, might have evolved a mechanism to remove the ubiquitination mark but still be cleared with autophagy through a different pathway. It was recently demonstrated that the *w*MelCS strain, but not the closely related *w*Mel, might have developed a trick to subvert the autophagy machinery by actively avoiding the ubiquitination in *D. melanogaster* hub cells (Deehan et al., 2021). These data show that even closely related *Wolbachia* strains exhibit stark differences in interaction with the host machinery, which might also be the case in our study. In addition, a few seemingly ubiquitinated bacteria were detected in systemic species as well as in PGCs of restricted species, suggesting that these single foci might be the false-positive signals derived from proteasomal degradation of proteins associated with ER (Meusser et al., 2005). As shown earlier, *Wolbachia* rely on host proteolysis to maintain a high titer within a cell and hence are closely interacting with the ER membranes (White et al., 2017). Therefore, we cannot dismiss the possibility that some of the interactions between ubiquitin and *Wolbachia* observed in the soma of WIL species are also false positive. However, the more pronounced pattern and higher number of co-localizations support the idea of bacterial tagging by ubiquitin for degradation in this RIT host.

The second step of bacterial elimination is characterized by ER membranes encircling the endosymbiont (**Figure 8B**). Various intracellular bacteria exhibit intimate contacts with the ER since it is a nutrient-rich organelle that is devoid of bactericidal effectors and thereby provides a safe niche for endosymbionts to survive and replicate (reviewed in Celli and Tsolis, 2015). As demonstrated in earlier studies, *Wolbachia* exerts close interaction with the ER membranes in different *D. melanogaster* tissues as well as in fly-derived cell lines (Voronin et al., 2004; Serbus et al., 2011; Strunov et al., 2016; White et al., 2017; Fattouh et al., 2019). Additionally, endosymbionts most likely receive the third outer membrane from the ER, which helps them to escape a cellular defence system (reviewed in Serbus et al., 2008). The ER, however, is not always a friendly environment for bacteria. Disruption of the secretory pathway by active endosymbiont interaction, causing ER stress, might lead to recognition by the innate immune system and cell defence response (reviewed in Celli and Tsolis, 2015). Moreover, the ER seems to provide a cradle for autophagosome formation (Hayashi-Nishino et al., 2009), which might ameliorate the elimination of bacteria. In our TEM studies we uncovered intimate interaction of rough ER membranes with *Wolbachia* in PAU embryos during the symbiont’s elimination process, which is in contrast to MEL embryos with rare and significantly lesser intimate contacts. Based on the results of our antibody staining against GABARAP, we speculate that ER membranes surrounding *Wolbachia* in PAU embryos serve as a scaffold for autophagosome formation. The role of ER membranes in the degeneration of bacteriocytes was also shown for the symbiotic Buchnera-Aphid system (Simonet et al., 2018). Additionally, ER encircling was recently demonstrated for damaged mitochondria elimination via mitophagy in mouse embryonic fibroblasts (Zachari et al., 2019). Very similar to our observation, not fully functional mitochondria are first ubiquitinated and then surrounded by ER strands, which provide a platform for mitophagosome formation and further degradation of the organelle. Given that mitochondria have alphaproteobacterial ancestry, both observations mentioned above strongly support our hypothesis of ER playing a key role in the somatic elimination of the *α*-proteobacteria *Wolbachia* in early RIT embryos by forming a cradle for autophagosome maturation.

The third step of bacterial elimination process is attraction of the autophagy machinery followed by autophagosome maturation (**Figure 8C**). It is known that autophagy plays an important role in defending the host cell against pathogens but in some cases the autophagy machinery can be hijacked by the intruder for its own survival (reviewed in Huang and Brumell, 2014). In systems with a mutualistic type of interaction, autophagy might be a key player in keeping the cost-benefit equilibrium in balance. Also for facultative symbiotic associations it was shown that *Wolbachia* density is regulated by autophagy (Voronin et al., 2012; Le Clec’h et al., 2012; Deehan et al., 2021). In our study, we observed *Wolbachia* accumulation mostly in PGCs during embryogenesis, whereas the rest of infection in the somatic part is being massively eliminated and subsequently restricted to certain isolated areas. Eventually, the adult flies exhibit highly abundant infection within the reproductive part of the gonad (nurse cells and oocyte) and restricted infection in somatic part, like follicle cells and nervous tissues. Such a specific tropism with a safe niche for bacteria in embryonic PGCs can be explained from the perspective of both symbiotic partners. On one hand, for ensuring their own maternal transmission, *Wolbachia* might specifically avoid autophagy in gonad precursors by actively blocking it with unknown effector proteins, which are released via type IV secretion system. As shown in the literature, some bacteria are able to counteract the host defence system by selectively preventing any of these three steps: detection, autophagy initiation or autophagosome formation (reviewed in Kimmey and Stallings, 2016; Wu and Li, 2019). This defence strategy of the symbiont also coincides with the downregulation of autophagy genes as observed in ovaries of the wasp *Asobara tabida* and the woodlouse *Armadillidium vulgare* (Kremer et al., 2012; Chevalier et al., 2012). Additionally, a recent study demonstrated that *w*MelCS strain of *Wolbachia* evolved a mechanism to subvert host autophagy in order to survive in hub cells and both *w*Mel and *w*MelCS are able to avoid elimination in the developing egg (Deehan et al., 2021). On the other hand, the PGCs themselves might lack extensive autophagic activity and thereby provide a safe environment for the *Wolbachia* to survive, replicate and being successfully transmitted via oocytes. In contrast to somatic cells, PGCs are transcriptionally quiescent during early embryonic stages (Cinalli et al., 2008) and activated only at later stages during their migration (Van Doren et al., 1998). It is conceivable that autophagy is blocked or impeded in germline stem cells during this quiescent state. Although, for this study, we did not conduct additional experiments to decipher the mechanism of preservation of bacterial infection in PGCs, it appears to be more plausible that the cell-specificity in development is a key regulator for *Wolbachia*’s fate. Thereby, during this critical step in early embryogenesis, PGCs are serving a safe haven for the maternally transmitted endosymbiont within the hostile somatic environment of massive autophagy in *Drosophila* species with the RIT phenotype.

The final step of the bacterial elimination process is degradation (**Figure 8D**). In our TEM studies we observed several abnormalities of *Wolbachia* morphology in the soma of PAU embryos during elimination of infection like stretching, bending and membrane vesiculation. Usually dying *Wolbachia* exhibit shrivelled, electron-dense structures surrounded by autophagosomal membranes (Wright and Barr, 1980; Min and Benzer, 1997; Zhukova and Kiseleva, 2012; Strunov et al., 2016), but the abnormalities observed in our study on RIT embryos are unique and represent a yet uncommon way of bacterial degradation. Although not found before with bacteria but associated with organelles, similarly stretched and bended structures were reported about stressed mitochondria in murine embryonic fibroblasts (Ding et al., 2012) and other mouse tissues (Gautam et al., 2019), linking these morphological deformations to autophagosome maturation by engulfing the cytoplasm and subsequent organelle degradation. In the latter more recent study, actual autophagosome formation was not confirmed by antibody staining but the authors speculated that mitochondria can undergo a self-destruction process, called mitoautophagy (Gautam et al., 2019). Morphologically similar ultrastructural abnormalities were also found with plastids of *Brassica napus* plants during the developmental switch from microspores to embryogenesis. Here, the authors experimentally verified these abnormal plastids with autophagosome formation and further elimination (Parra-Vega et al., 2015). Taken together, the discovered deformities of *Wolbachia* morphology in embryogenesis of RIT *Drosophila* hosts most likely represent the first report of a non-canonical degradation process of bacteria through autophagy, which was only found in organelles before.

## Conclusion

In the present study we reconstructed the mechanism of restricting *Wolbachia* infection by autophagy in three different neotropical *Drosophila* species. These data present a unique way of symbiont density regulation by the host during a specific period in embryogenesis, which coincides with maternal-to-zygote transition. They also demonstrate how the cost-benefit equilibrium between the host and the symbiont is maintained long-term by keeping a safe niche in the reproductive part, thereby being transmitted to the next generation, while being eliminated from most of the somatic part to reduce potential costs. It is still unclear how *Wolbachia* escapes elimination in PGCs and in the soma of systemic species. One possibility is a unique marker on the bacterial surface, which is specifically recognized by a native host but further transinfection experiments with various *Wolbachia* strains into different *Drosophila* backgrounds might give us the answers.

## Acknowledgements

We thank Matthias Schäfer and Silvia Bulgheresi for critical and constructive discussions during the whole period of the project and for providing the antibodies Matthias Schäfer (Medical University of Vienna), the groups of Fumiyo Ikeda (IMBA) and Sasha Marten (MFPL), as well as Thomas Hummel (University of Vienna) and Anne Ephrussi (EMBL). We thank Ivanna Fedorenko from Katy Schmidt’s group for substantial and excellent technical support. We are also very thankful to Aurélie Hua-Van for fly sampling in French Guiana, France, and the Nouragues research field station (managed by CNRS), which benefits from “Investissement d’Avenir” grants managed by Agence Nationale de la Recherche (AnaEE France ANR-11-INBS-0001; Labex CEBA ANR-10-LABX-25-01) to WMJ. The project was funded by the Austrian Science Fund FWF grants P28255-B22 to WJM and P32275 to MK

## Materials and methods

### Fly stocks and husbandry

Seven different species from four *Drosophila* subgroups were used in this study: *D. melanogaster* (MEL) and *D. simulans* (melanogaster subgroup), *D. paulistorum* (PAU), *D. willistoni* (WIL), *D. tropicalis* (TRO) (willistoni subgroup), *D. septentriosaltans* (SPT) (saltans subgroup) and *D. sturtevanti* (STV) (sturtevanti subgoup). All the species mentioned above were naturally infected with specific *Wolbachia* strain (*w*Mel, *w*Au, *w*Pau, *w*Wil, *w*Tro, *w*Spt and *w*Stv, respectively). Additionally, the stably transinfected *w*Wil/STC line was used in the experiment, generated in 2006 by injecting *w*Wil *Wolbachia* from *D. willistoni* into *D. simulans* STC early embryos, which were cleared from the native *w*Au *Wolbachia* with antibiotics. For more details on flies used in the study see the **Table 1**. All lines were kept at 22–25°C on a 12 h light-dark cycle and fed a typical molasses, yeasts, cornmeal and agar food.

### RNA-DNA fluorescent in situ hybridization

Tissues (adult brains, larval CNS, adult ovaries, larval ovaries) from at least ten females per *Drosophila* species/line were dissected in ice-cold RNase-free 1x phosphate buffered saline (PBS) and fixed in 3.7% formaldehyde in RNase-free PBS for 15-20 min at room temperature and consequently washed 3 times 5 min each with PBTX (1xPBS, 0.3% Triton-X 100). Embryos from listed *Drosophila* species were collected and fixed according to a standard protocol (Rothwell and Sullivan et al., 2007).

All fixed samples were hydrated in prewarmed 4xSSC buffer with 10% formamide and hybridized at 37 °C overnight in the same buffer containing 10% of dextran sulfate and 0.5 nmol of W1/W2 probes specifically targeting *Wolbachia* 16S rRNA (Heddi et al., 1999) labeled with Oregon Green (488) or Texas Red (596) fluorophore. Samples were then washed twice for 30 min at 37 °C in prewarmed 4xSSC buffer with 10% formamide. For preparation of larval CNS, ovaries and adult ovaries, tissues were additionally incubated in Alexa Fluor 488 phalloidin (Invitrogen, USA; 1:100 dilution in 1xPBS) for 1h at room temperature to stain F-actin. Finally, after washing samples 2 times with 1x PBS, they were mounted in Roti®-Mount FluorCare with DAPI (Carl Roth, Germany) on microscope slides.

Samples were analyzed on Olympus FluoView FV3000 confocal microscope. Beam paths were adjusted to excitation/emission peaks of used fluorophores: 569/591 nm for CAL Fluor Red 590 (*Wolbachia*), 488 nm for phalloidin and 350/450 nm for 4′,6-diamidin-2-phenylindol (DAPI).

### FISH combined with immunofluorescence (FISH/IF)

For combination of FISH with antibody staining we first conducted *in situ* hybridization as described in the section above. After washing steps in prewarmed 4xSSC buffer samples were incubated in 5% bovine serum albumin (BSA) for 1h at room temperature constantly shaking. Then they were washed once with 1%BSA and incubated with a primary antibody (diluted in 1xPBTX with 1%BSA) overnight at 4°C constantly shaking. The following day the samples were washed 3 times 10 min each in 1xPBTX and incubated in a secondary antibody (diluted in 1xPBTX with 1%BSA) for 1h at room temperature constantly shaking. After washing 3 times 10 min each with 1xPBTX samples were stained with Alexa Fluor 488 phalloidin (Invitrogen, USA; 1:100 dilution in 1xPBS). Then they were washed 2 times with 1x PBS and mounted in Roti®-Mount FluorCare with DAPI (Carl Roth, Germany) on microscope slides

### Antibodies

The following primary antibodies were used in this study: anti-Deadpan (guinea pig, polyclonal; 1:1000; Eroglu et al., 2014), anti-Asense (guinea pig, polyclonal; 1:100; Eroglu et al., 2014), anti-Repo (rabbit, polyclonal; 1:1000; gift of G. Technau), anti-Vasa (rat, polyclonal; 1:500; gift of A. Ephrussi), anti-GABARAP (rabbit, polyclonal; 1:200; gift of S. Martens), anti-FK2 (mouse, monoclonal; 1:200; gift of F. Ikeda), anti-GRP78/BiP (rabbit, polyclonal; 1:500; Abcam, Cambridge, UK). The following secondary antibodies were used in this study: goat anti-mouse Alexa Fluor 488 (1:500), goat anti-mouse Cy5 (1:500), goat anti-rabbit Alexa Fluor 488 (1:500), goat anti-guinea pig Cy3 (1:500), goat anti-rat Alexa Fluor 488 (1:500). All secondary antibodies were obtained from Invitrogen, USA.

### Transmission electron microscopy

*Drosophila* embryos were collected the same way as for FISH and then fixed in 2.5% (w/v) glutaraldehyde in 0.1 M sodium cacodylate buffer (pH 7.2) for 2.5 h. This was followed by three washes in the same buffer for 5 min each and post-fixation in 1% (w/v) OsO_4_ and 0.8% (w/v) potassium ferrocyanide for 1 h. Samples were then placed in a 1% aqueous solution of uranyl acetate (Serva, Heidelberg, Germany) for 12 h at 4°C and dehydrated in an ethanol series (30%, 50%, 70%, 96% for 10 min, and 100% for 20 min) and acetone (twice, for 20 min). Ultra-thin sections of embedded samples (Agar 100 Resin; Agar Scientific Ltd., Essex, UK) were obtained with a Reichert-Jung ultracut microtome, stained with Reynolds lead citrate and examined in an FEI Tecnai 20 electron microscope (FEI Eindhoven, Netherlands) equipped with 4K Eagle CCD camera. Images were processed with Adobe Photoshop.

### Analysis and quantification of Wolbachia localization in the tissue

Calculation of restriction index (RI), aggregation of infection and *Wolbachia* density:

We define a restriction index (RI) to quantify the pattern of *Wolbachia* localization as number of uninfected cells divided by total number of cells:

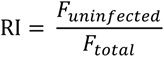

*F_uninfected_* and *F_total_* in adult brains and larval CNS was calculated by superimposing a grid (25x25µm) on the whole tissue image in Photoshop CS6 and quantifying the number of uninfected and total number of grids containing the tissue. The RI value varied from 0 (no restriction) to 1 (full restriction). In total, 10 samples per each *Drosophila* species and each tissue were analyzed (more than 1200 grid cells for adult brains and approximately 400 grid cells for larval nervous tissues of each species).

The RI of infection in adult and larval ovaries was calculated by dividing the number of uninfected follicle cells from a central section of egg chamber (for the former) or somatic cells related to terminal filament (for the latter) to the total number of cells analyzed. In total, 10 samples per each *Drosophila* species and each tissue were analyzed (more than 400 cells for adult ovaries and more than 170 cells for larval ovaries of each species). The RI of infection in somatic cells around primordial germ cells (PGCs) in embryos was quantified by drawing a 50x50μm square around PGCs, counting the number of uninfected cells within this square and dividing it to the total number of cells. In total, 10 samples per each *Drosophila* species and each tissue were analyzed (more than 300 cells for each species).

The RI of infection in neuroblasts of embryonic head was quantified by counting the number of uninfected cells (stained with anti-Deadpan antibody specific to neuroblasts) and dividing it to the total number of neuroblasts. In total, 10 samples per each *Drosophila* species and each tissue were analyzed (more than 400 neuroblasts for each species).

Aggregation of *Wolbachia* in larval CNS was calculated by quantifying the average number of infected neighboring cells forming a cluster in each tissue. In total, 8 samples per each *Drosophila* species were analyzed (61-65 cell clusters for SIT, 26-32 cell clusters for RIT and 56 cell clusters for transinfected line).

*Wolbachia* density within a neuroblast of larval CNS, within an egg chamber of an ovary or an embryo was quantified with Fiji (Schindelin et al., 2012) by measuring the area of bacterial signal within the region of interest (ROI) and dividing it to the total area of the ROI. In total, at least 5-10 samples per each *Drosophila* species and each tissue were analyzed. The detailed description of this procedure can be found in Strunov et al., 2017.

### Statistics

All statistical analyses were carried out using *R* version 3.3.2 (R-Core Team, 2020). For *Wolbachia* distribution in adult and larval brains and ovaries we analyzed the count data based on generalized linear models (GLM) with a Poisson error structure. To test for significance of a given predictor variable, we compared the full model including all factors to a reduced model excluding the given factor by analysis of deviance with *χ*^2^ tests using the *R* function *anova*. For the rest of the data, we assume that the data is normally distributed and calculated one-way ANOVAs. We further applied post-hoc Tukey HSD test to test for significant difference among factor levels using the R function *TukeyHSD*.

## Supplemental Figures

**Figure S1.**
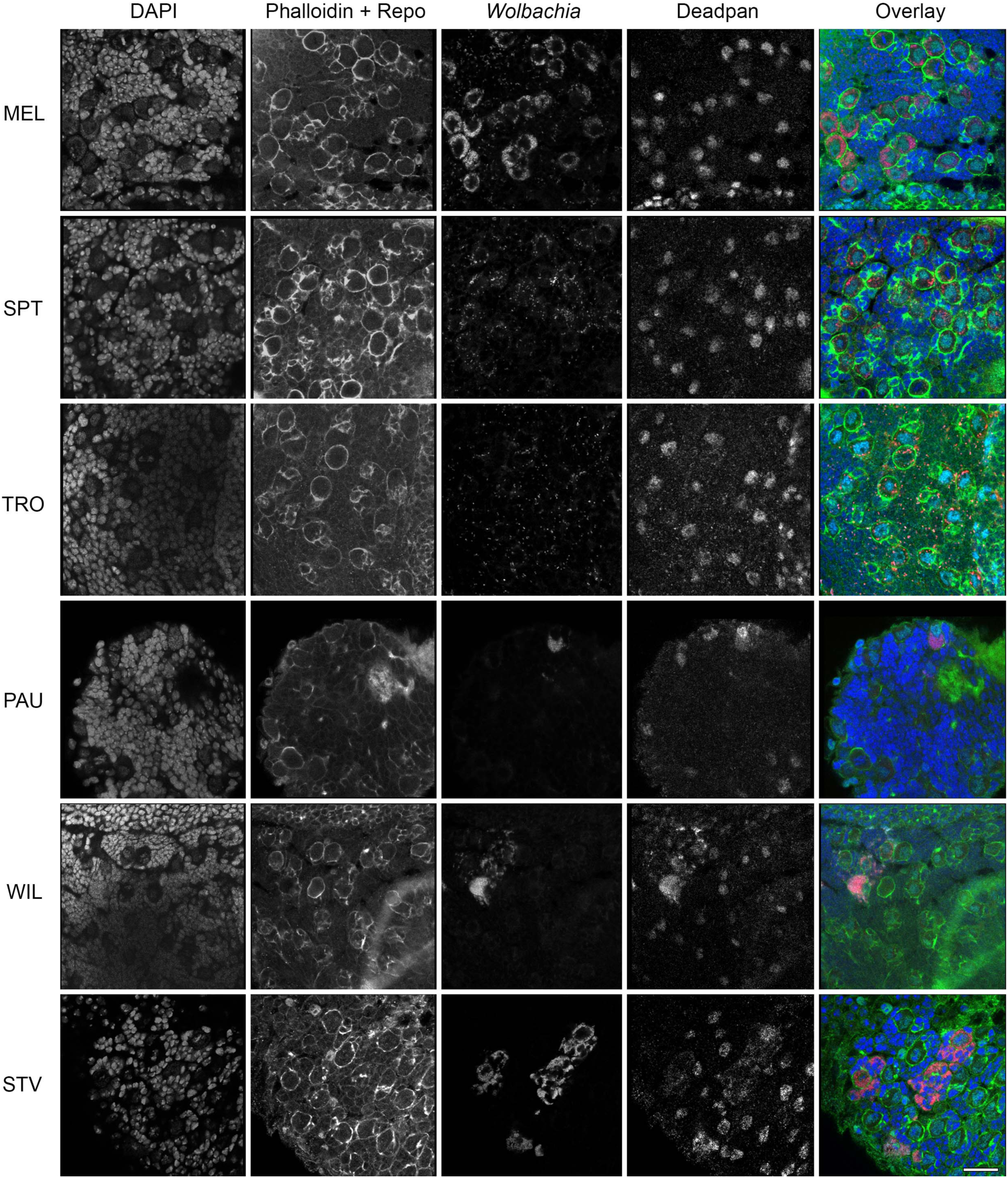
*Wolbachia* infection in neuroblasts of the CNS of 3^rd^ instar *Drosophila* larvae. Sequential RNA-FISH using *Wolbachia*-specific 16S rRNA probe (red) followed by immunofluorescent staining with anti-Repo (glial cells, green) and anti-Deadpan (neuroblasts, cyan) antibodies of 3^rd^ instar larval CNS. DNA is stained with DAPI (blue), and actin with Phalloidin (green). For each *Drosophila* species 10 organs were analyzed. Scale bar: 20 μm.

**Figure S2.**
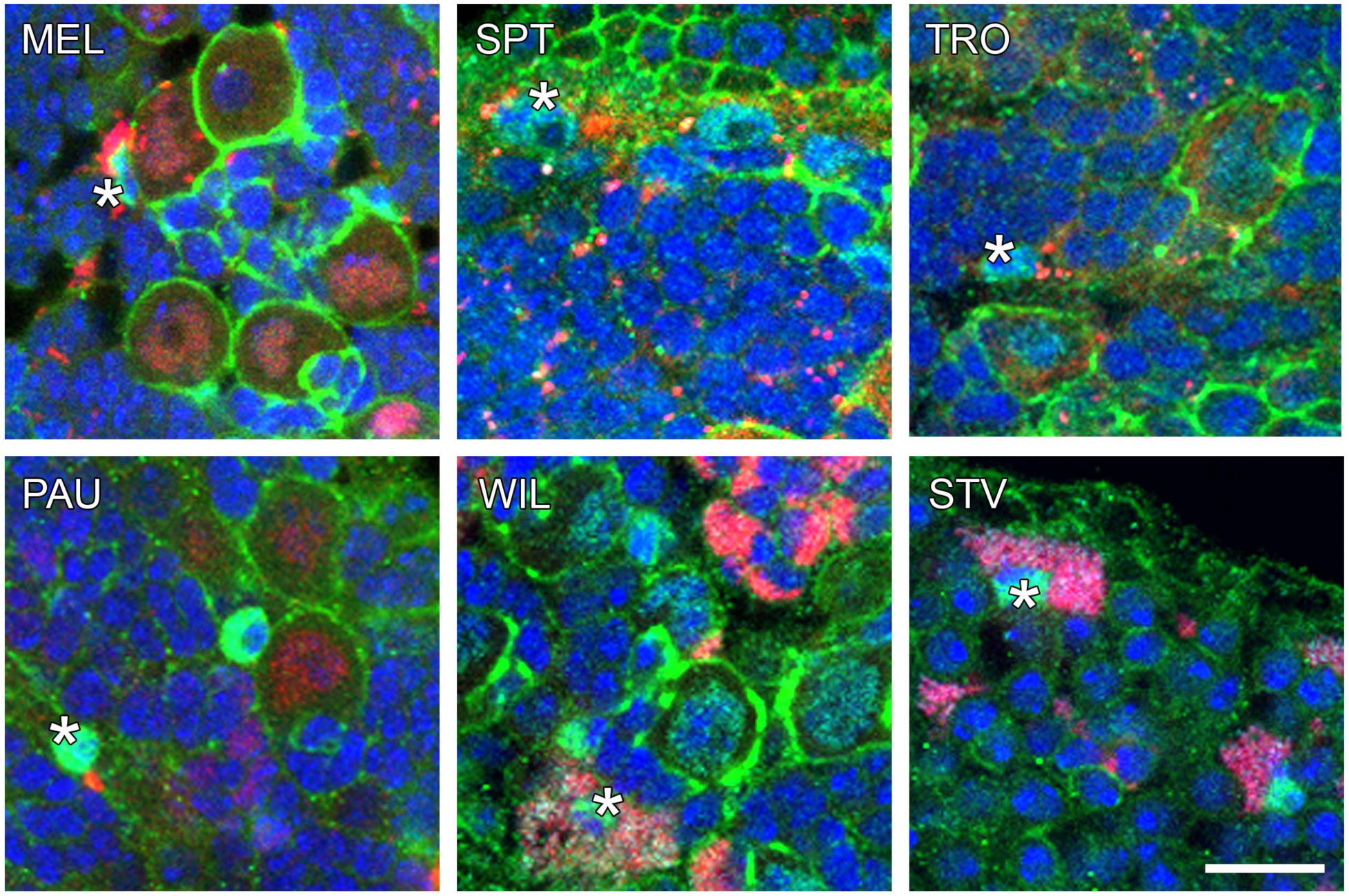
*Wolbachia* infection of glial cells in the CNS of 3^rd^ instar *Drosophila* larvae. Sequential FISH using *Wolbachia*-specific 16S rRNA probe (red) followed by immunofluorescent staining with anti-Repo (glial cells, green) and anti-Deadpan (neuroblasts, cyan). DNA is stained with DAPI (blue), and actin with Phalloidin (green). Asterisks indicate a glial cell infected with *Wolbachia*. For each *Drosophila* species 10 organs were analyzed. Scale bar: 10 μm.

**Figure S3.**
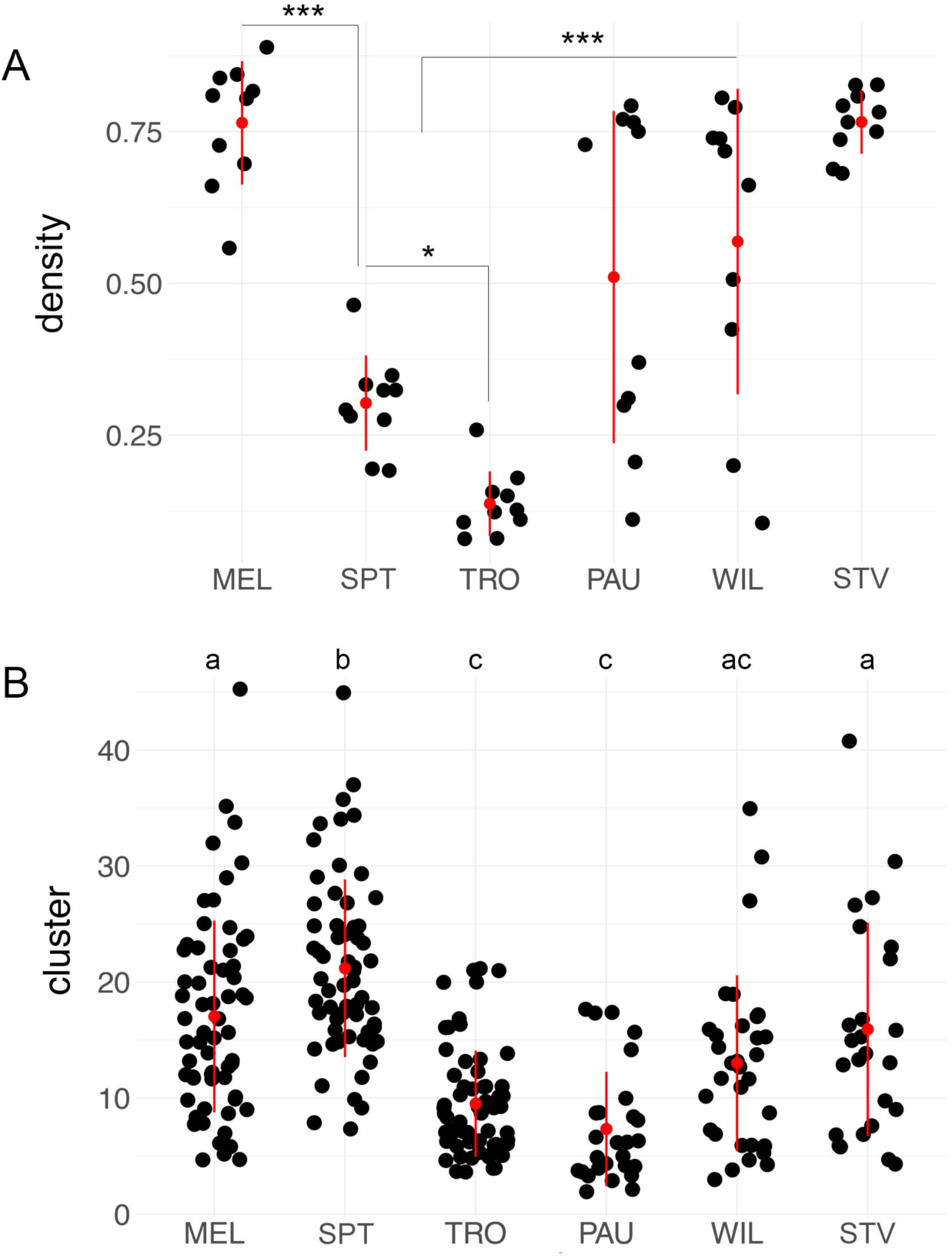
Density and cluster aggregation of *Wolbachia* in the CNS of 3^rd^ instar *Drosophila* larvae. (**A**) Density within 10 neuroblasts of 3 individual brains was quantified with Fiji as a bacterial load area divided by an area of cell cytoplasm. (**B**) Aggregation of infection in the larval CNS of six *Drosophila* species was analyzed from bacterial clusters in 5 individual brains (61-65 clusters for SIT and 26-32 clusters for RIT) by quantifying the number of neighboring infected neurons in groups. In **A** asterisks denote statistical significance (*, *p*<0.05; ***, *p*<0.001; One-way ANOVA with Tukey HSD Test). In **B** statistical significance is shown with letters (*p*<0.05, One-way ANOVA with Tukey HSD Test). Red bars show standard deviation, red dots designate the mean value. For more details, see Supplemental data file.

**Figure S4.**
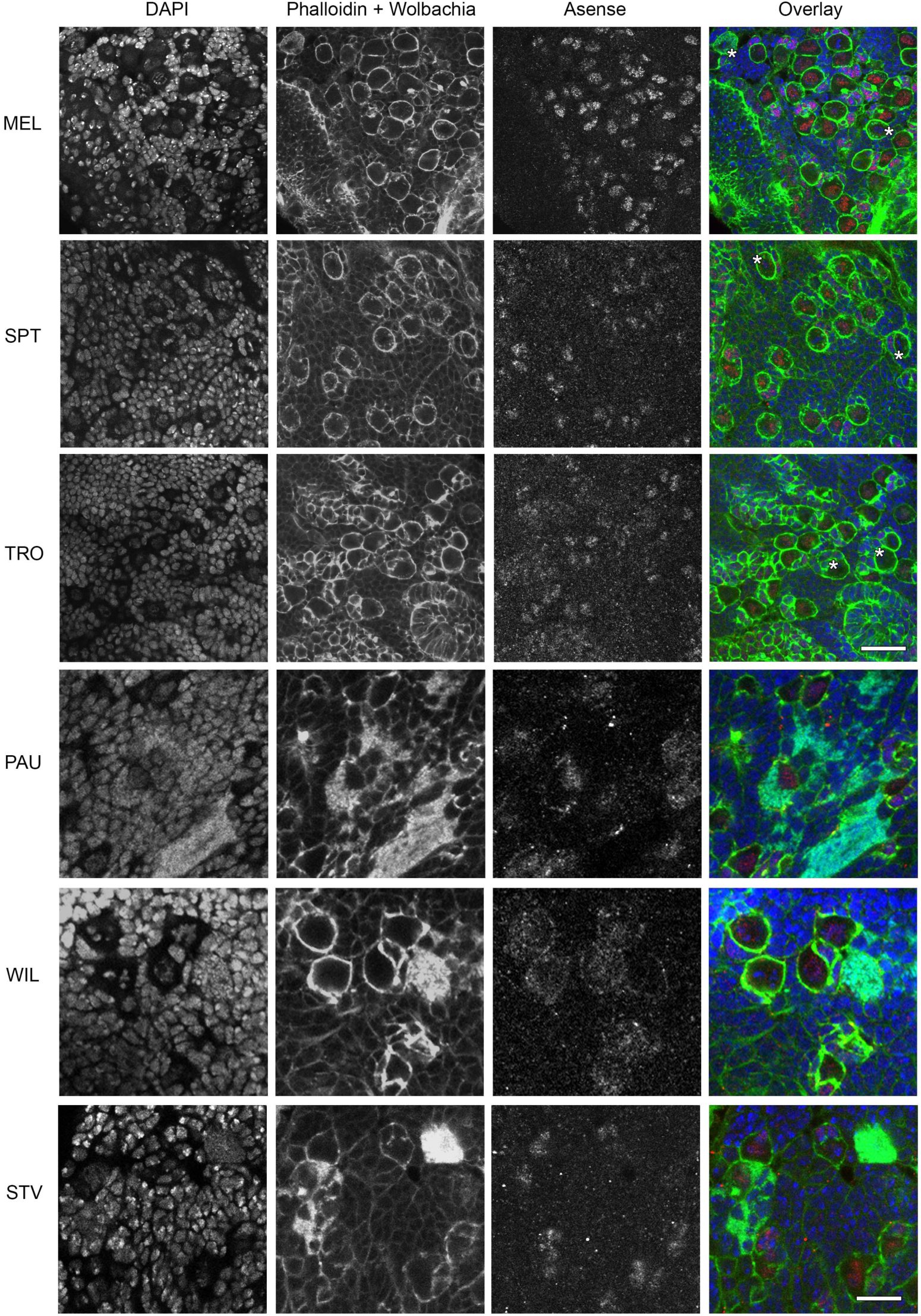
*Wolbachia* infection in type I and II neuroblasts in the CNS of 3^rd^ instar *Drosophila* larvae. Sequential FISH using *Wolbachia*-specific 16S rRNA probe (green dots) and immunofluorescent staining with anti-Asense antibody (red), which is diagnostic for Type I neuroblasts, of 3^rd^ instar larval CNS. DNA is stained with DAPI (blue), actin with Phalloidin (green). Asterisks depict type II neuroblasts, which are Asense-negative, infected with *Wolbachia* (green dots). In total 10 brains were analyzed for each species. Scale bar: 20 μm (MEL, SPT, TRO), 10 μm (PAU, WIL, STV).

**Figure S5.**
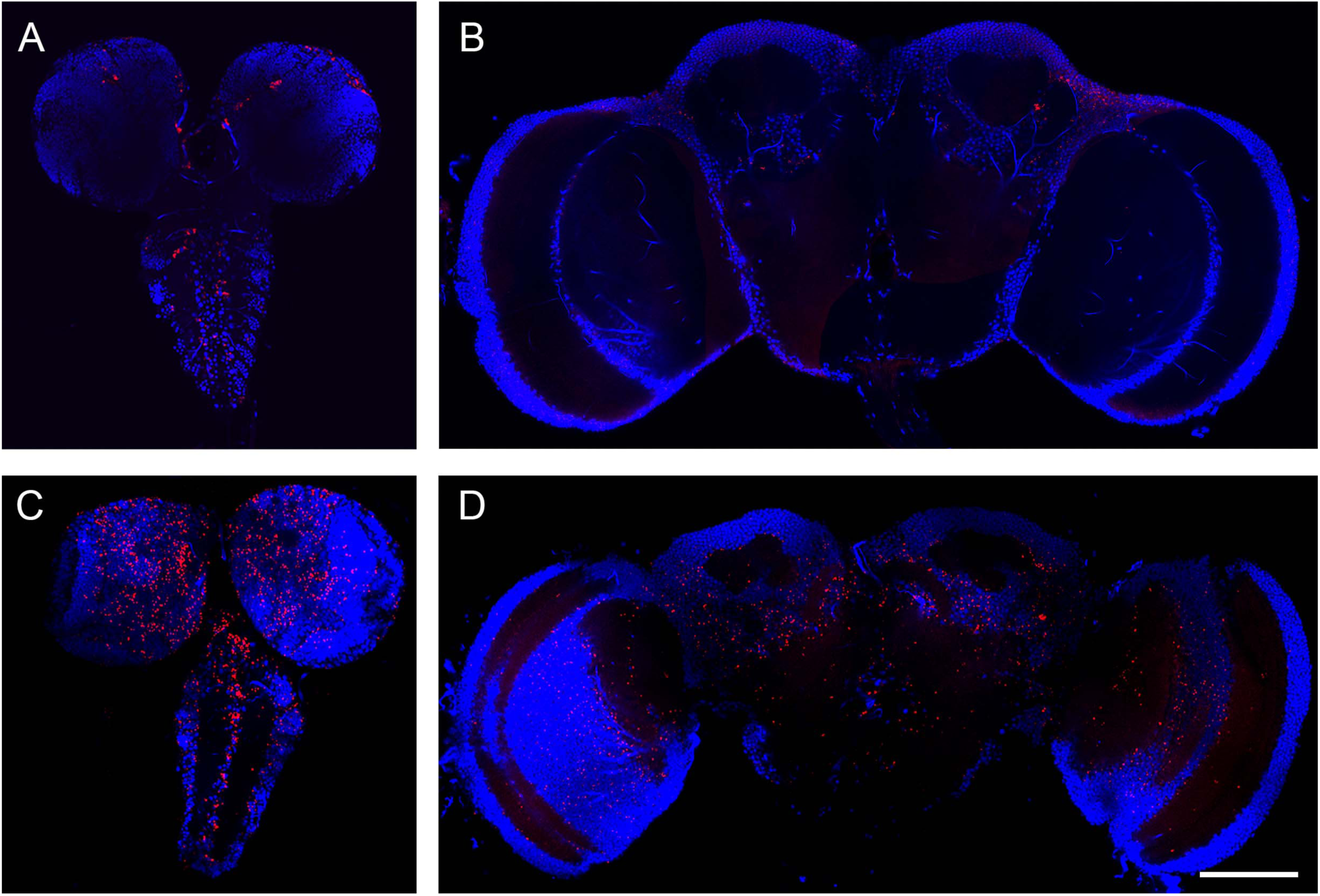
*Wolbachia* infection in nervous tissues of *Drosophila lehrmanae* (A, B) from sturtevanti subgroup and *Drosophila prosaltans* (C, D) from saltans subgroup. Fluorescent *in situ* hybridization on 3^rd^ instar larval CNS (**A, C**) and adult brains (**B, D**) using 16S rRNA *Wolbachia*-specific probe (red). DNA is stained with DAPI (blue). Note restriction of *Wolbachia* in *D. lehrmanae* and systemic infection in *D. prosaltans*. Scale bar: 50 μm.

**Figure S6.**
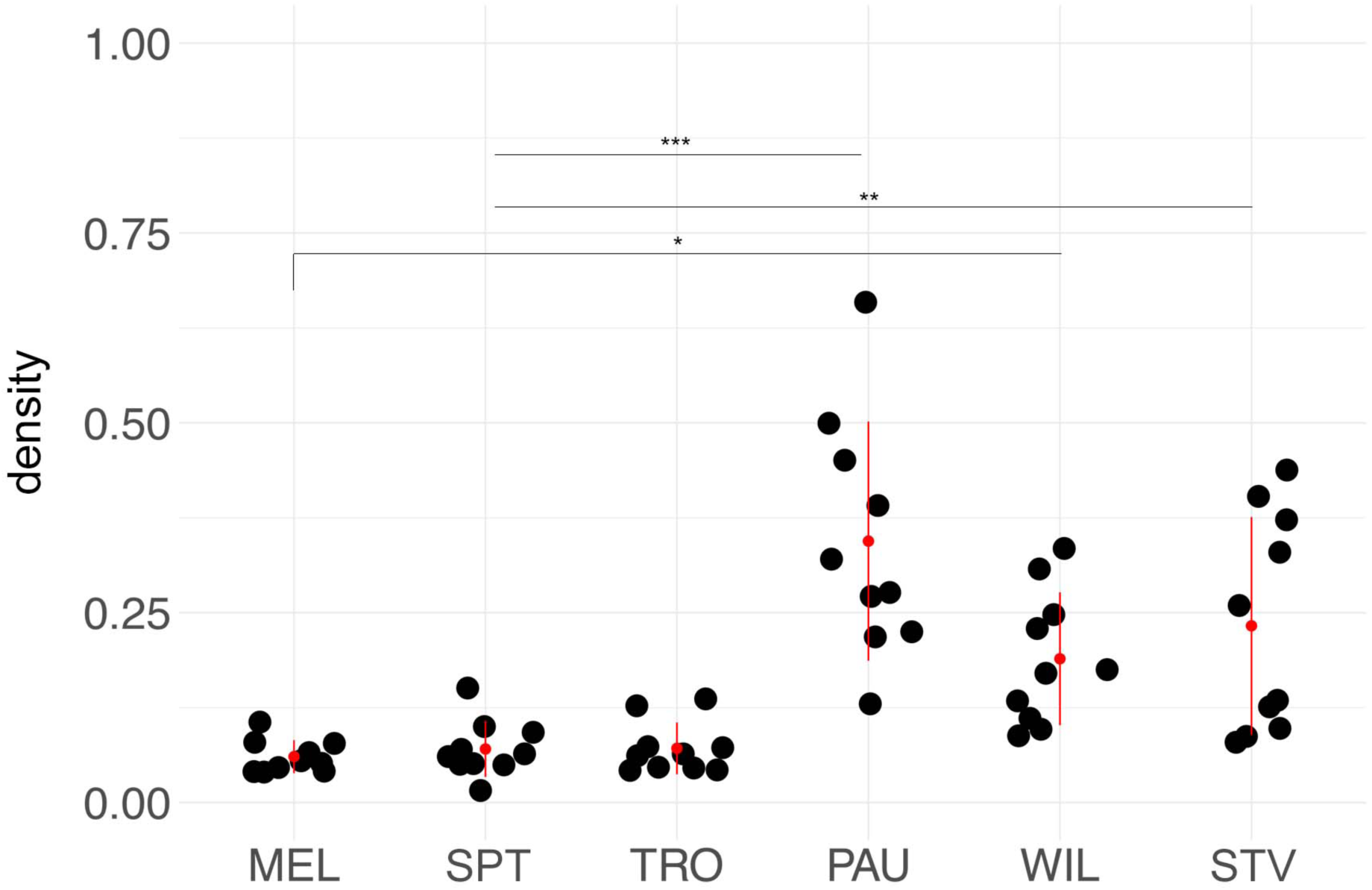
*Wolbachia* densities in the nurse cells of stage 3-5 ovaries of neotropical *Drosophila* species. The bacterial density was analyzed in all six *Drosophila* species with Fiji as bacterial infection area in an egg chamber divided by an area of the chamber. Asterisks denote statistical significance (*, *p*<0.05; **, *p*<0.01; ***, *p*<0.001). Red bars show standard deviation, red dots designate the mean value. In total, ten egg chambers were analyzed for every species (Supplemental data file).

**Figure S7.**
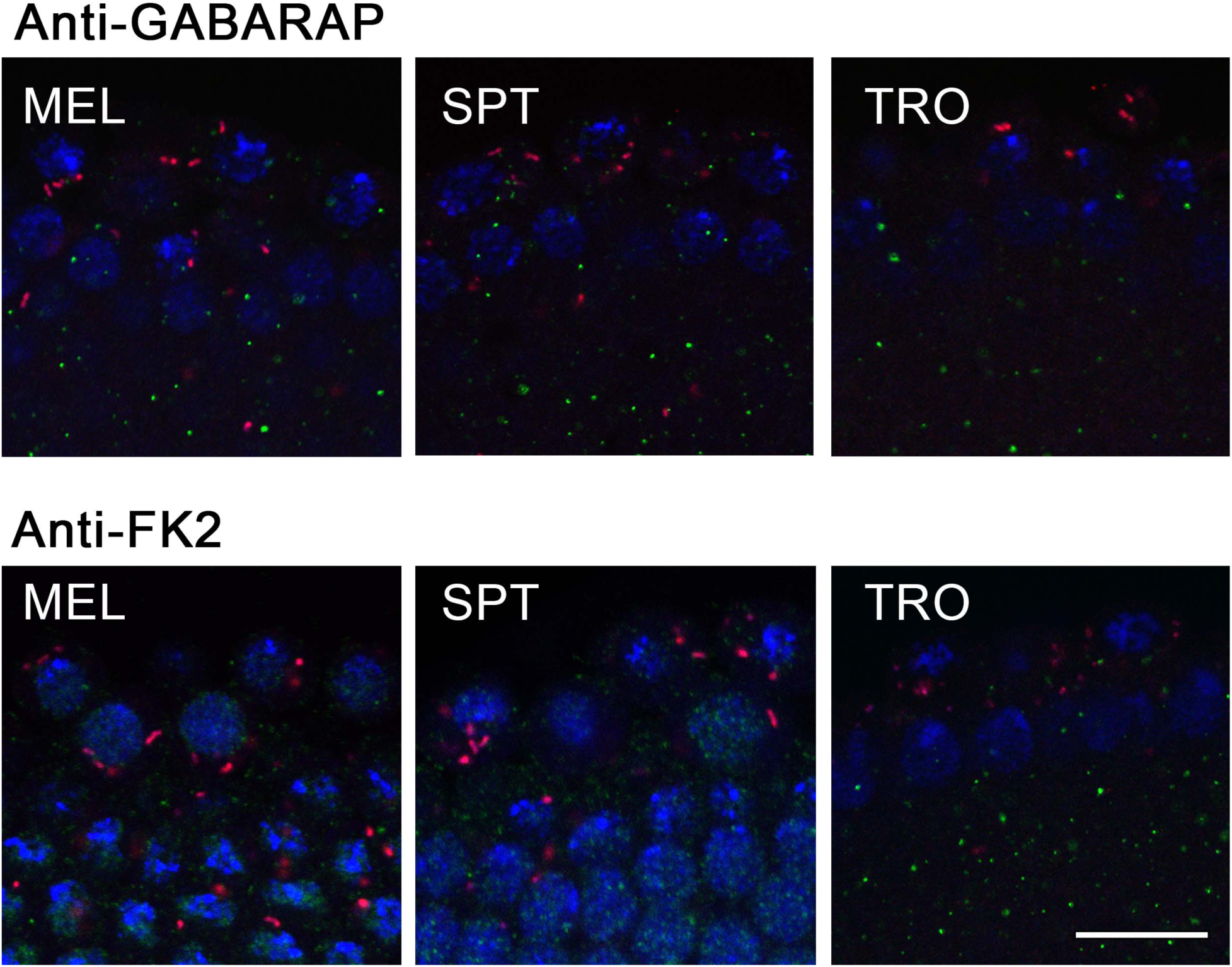
Absence of autophagy and ubiquitination of *Wolbachia* in primordial germ cells of *Drosophila* with SIT pattern of infection at stage 5 of embryogenesis. Sequential FISH using *Wolbachia*-specific 16S rRNA probe (red) and immunofluorescent staining with the two autophagy-specific antibodies, i.e., anti-GABARAP (green, upper panel) and anti-FK2 (green, lower panel) on PGCs of embryos from the six species at the cellularization stage. DNA is stained with DAPI in blue. For each *Drosophila* species five embryos were analyzed. Scale bar: 20 μm.

**Figure S8.**
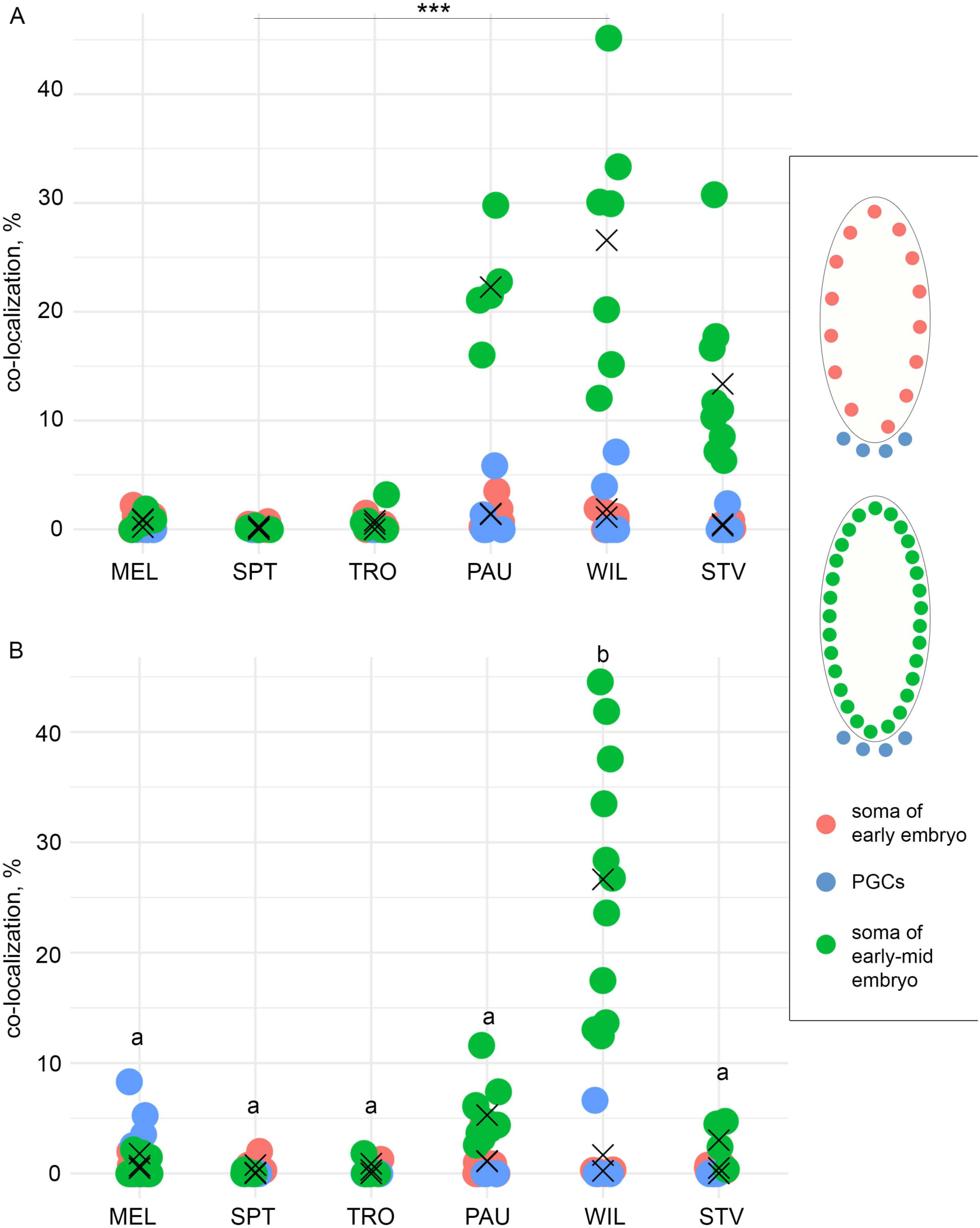
*Wolbachia* co-localization with anti-GABARAP (A) and anti-FK2 (B) antibodies. Co-localization was assessed using JACoP Fiji plugin. Each dot represents percentage of co-localization in a single embryo in the soma at stages 3-4 and stages 5-6 or PGCs of both (stages were fused due to the absence of differences). On **A** asterisks denote statistical significance only for soma of early-mid embryo at stages 5-6 (***, *p*<0.001; One-way ANOVA with Tukey HSD Test). On **B** letters indicate statistical significance only for soma of early-mid embryo at stages 5-6 (*p*<0.001, One-way ANOVA with Tukey HSD Test). “X” symbol demonstrates the mean value. For every species and every stage 4-11 embryos were analyzed (Supplemental data file).

**Figure S9.**
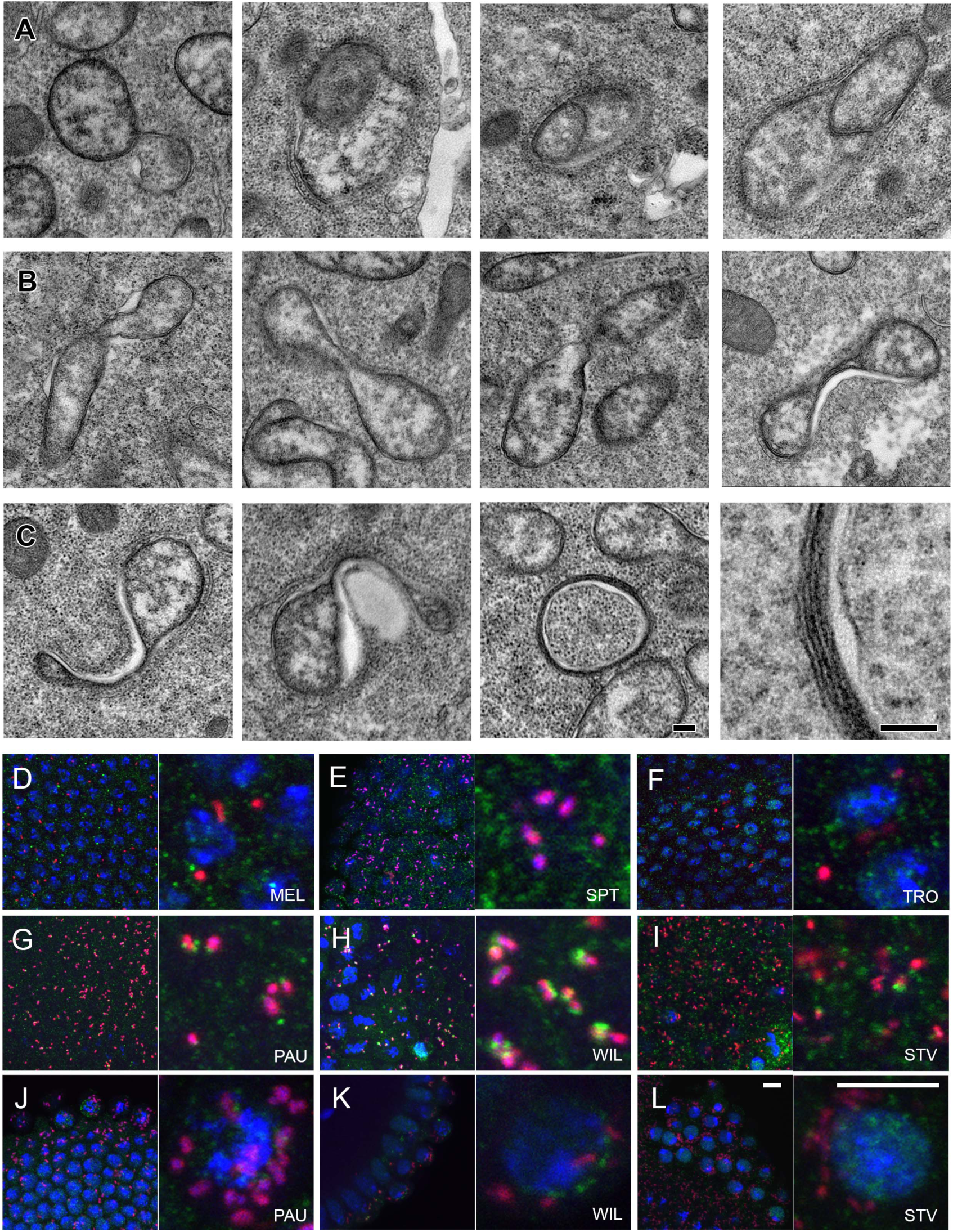
*Wolbachia* interactions with the host cell. Transmission electron microscopy images of abnormal *Wolbachia* in early gastrulating PAU embryos in the soma (**A-C**) demonstrating abnormalities in morphology like vesicle formation (**A**), stretching (**B**) and membrane extrusions (**C**). Sequential FISH using *Wolbachia*-specific 16S rRNA probe (red) and immunofluorescent staining with anti-FK2 (green) antibody (**D-L**). Note the absence of ubiquitination in SIT species (**D-F**) and co-localization of anti-FK2 with *Wolbachia* in RIT species (**G-I**). Also note the absence of co-localization of anti-FK2 with bacteria in PGCs of restricting species (**J-L**). Scale bar: 0.1 μm (**A-C**), 10 μm (**D-L**).

**Figure S10.**
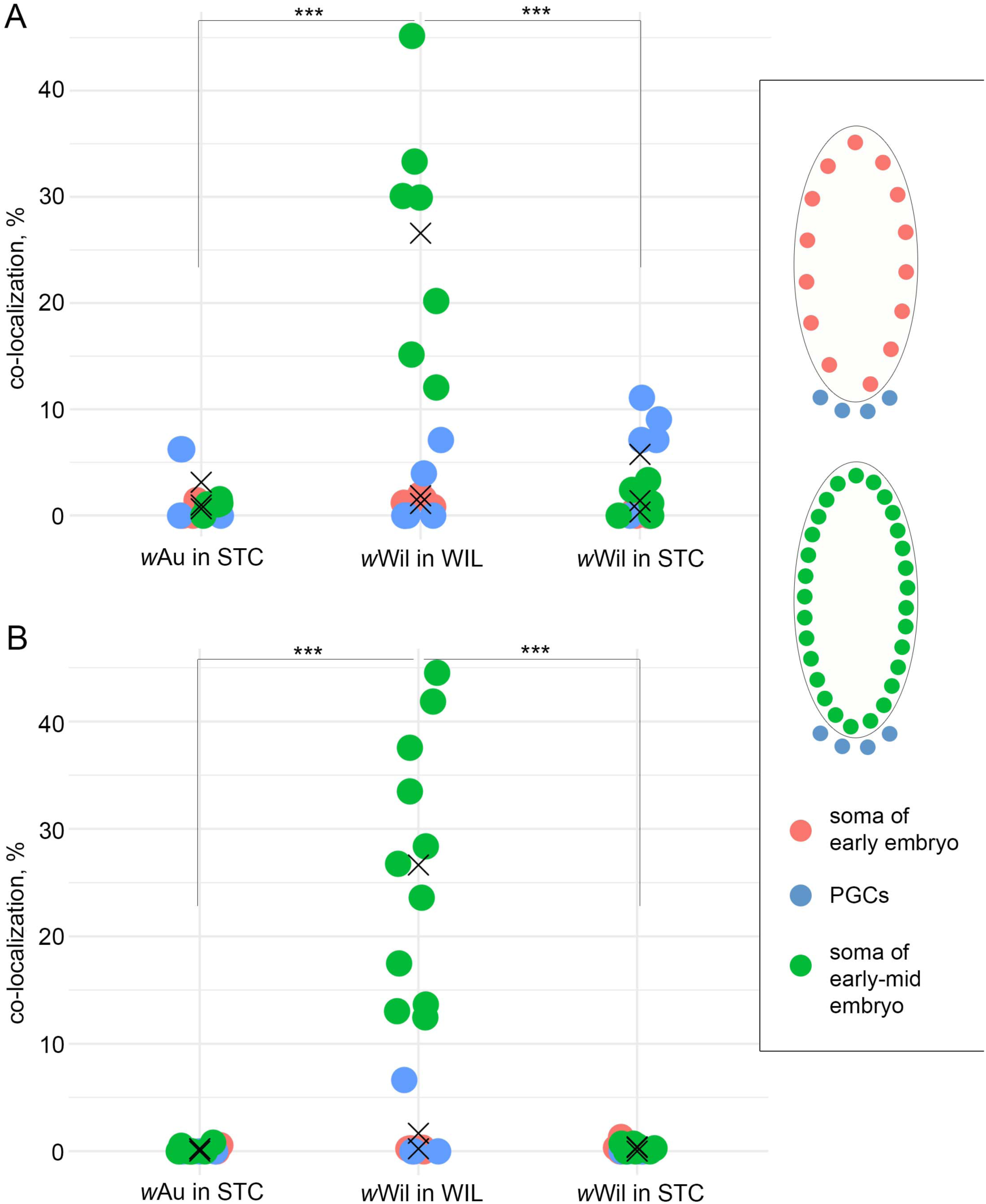
*Wolbachia* co-localization with anti-GABARAP (A) and anti-FK2 (B) antibodies. Co-localization was assessed using JACoP Fiji plugin. Each dot represents percentage of co-localization in a single embryo in the soma at stage 5 and stage 6 or PGCs at both stages (stages were fused due to the absence of differences). Asterisks denote statistical significance only for soma of early-mid embryo at stage 6 (***, *p*<0.001; One-way ANOVA with Tukey HSD Test). “X” symbol demonstrates the mean value. For every species and every stage 4-11 embryos were analyzed (Supplemental data file).

**Figure S11.**
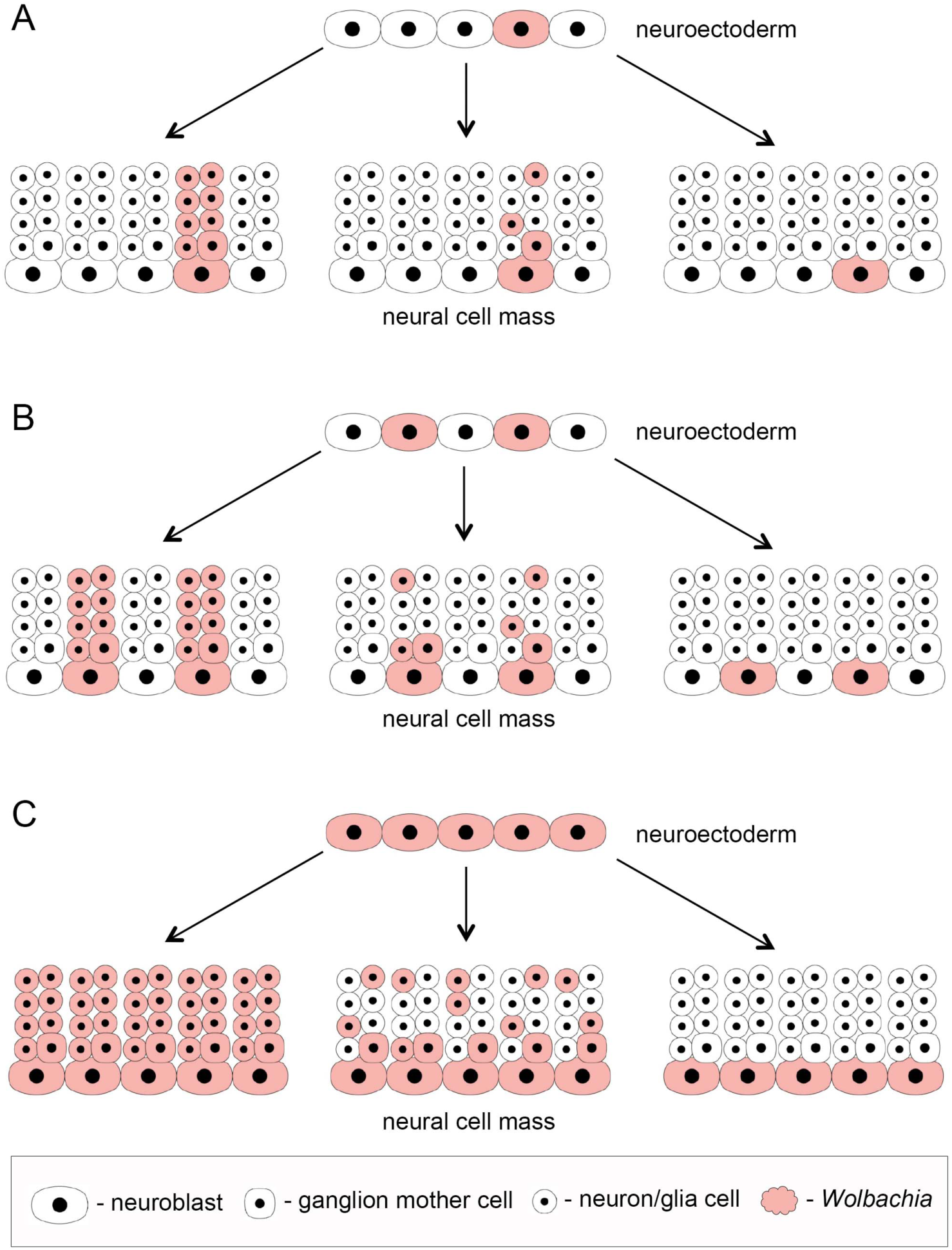
Description of all possible variants of *Wolbachia* distribution patterns during fly development exemplified on the central nervous system formation. The scheme demonstrates *Wolbachia* dissemination efficiency during mitosis of neuroblasts from the neuroectoderm with different starting numbers of infected stem cells (niches) – low (A), moderate (B) and high (C). Each neural cell mass picture demonstrates the percentage of cells in the progeny of a single neuroblast receiving the infection.

**Table S1.** The role of bacterial and host factors in regulating the distribution and density of the infection.

| references | tissue | tropism regulated by | titer regulated by |
| --- | --- | --- | --- |
| Veneti et al., 2004 | ovaries and testes | *Wolbachia* | host |
| Serbus and Sullivan, 2007 | ovaries | *Wolbachia* | host |
| Albertson et al., 2013 | CNS | *Wolbachia* | *Wolbachia* + host |
| Toomey et al., 2013 | ovaries | *Wolbachia* | NA |
| Toomey, Frydman, 2014 | testes | *Wolbachia* + host | NA |
| this study | CNS + ovaries | host | *Wolbachia* |

